# Why are G-quadruplexes good at preventing protein aggregation?

**DOI:** 10.1101/2022.08.23.504968

**Authors:** Theodore J. Litberg, Rajesh Kumar Reddy Sannapureddi, Zijue Huang, Ahyun Son, Bharathwaj Sathyamoorthy, Scott Horowitz

## Abstract

Maintaining a healthy protein folding environment is essential for cellular function. Recently, we found that nucleic acids, and G-quadruplexes in particular, are potent chaperones for preventing protein aggregation. With the aid of structure-function and NMR analyses of two G-quadruplex forming sequences, PARP-I and LTR-III, we uncovered several contributing factors that affect G-quadruplexes in preventing protein aggregation. Notably, three factors emerged as vital in determining holdase activity of G-quadruplexes: their structural topology, structural dynamics, and oligomerization state. These factors together appear to largely dictate whether a G-quadruplex is able to prevent partially misfolded proteins from aggregating. Understanding the genesis of G-quadruplexes’ power as chaperones is an important facet to elucidating various protein aggregation diseases.

**Key Points:**

- How nucleic acids act as protein chaperones is currently unknown.
- G-quadruplexes are excellent at preventing protein aggregation, and here we describe basic tenets of this activity.
- This activity could help design treatments for multiple neurodegenerative diseases.

## Introduction

Molecular chaperones are molecules (often proteins) that help determine the destiny of their nascent, misfolded, or aggregated protein clients, whether it be refolding or degradation (1). It has recently been shown that nucleic acids can also be potent molecular chaperones, preventing protein aggregation (also called holdase activity) on a per-weight basis more effectively than any known chaperone protein (2). Although bulk DNA can have holdase properties, further investigation into the structural and sequence dependence of chaperone activity found that G-quadruplexes possess powerful holdases activity *in vitro* and are able to improve the folding environment of *E. coli* (3).

G-quadruplexes are a secondary structure in guanine-rich DNA or RNA, where four guanines form a square plane stabilized by a metal cation (often potassium or sodium). These planes can then stack to form an intramolecular G-quadruplex within the same polynucleotide chain, or intermolecular G-quadruplexes with other chains. G-quadruplexes can also adopt a variety of different topologies based on the directionality of their backbone: parallel, hybrid 3+1, anti-parallel 2+2, and anti-parallel (4). (Figure 1) We previously found that the holdase activity of a G-quadruplex is loosely correlated to topology, with mixed hybrid 3+1 being the most active, followed by parallel, while anti-parallel G-quadruplexes display negligible holdase activity (3). Having shown that G-quadruplexes are potent holdases and with their activity dependent on structural topology, we sought to investigate the underlying causes of this activity.

**Figure 1.**
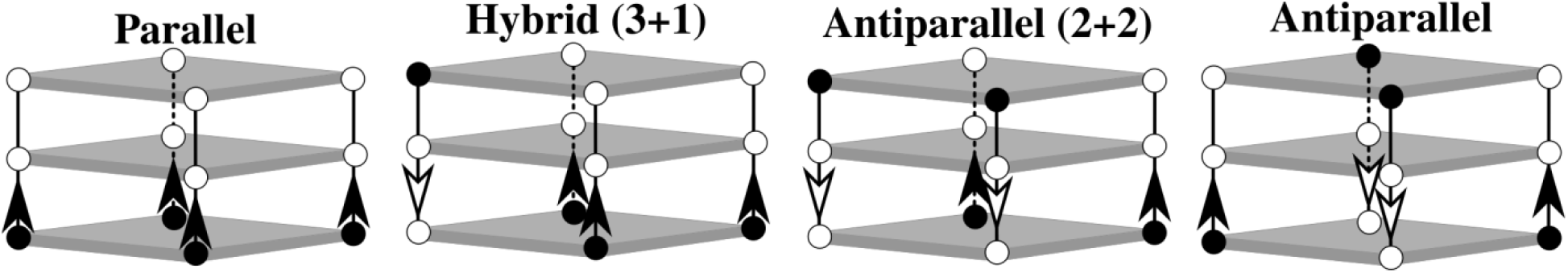
G-quadruplex topologies based on strand arrangement. Each tetrad is represented as grey box. Each strand is shown with arrows indicating directionality (5’ to 3’), each circle represent deoxyribose sugar and phosphate group. 5’ is shown as filled (black) circle.

Additionally, G-quadruplexes play important roles in neurodegenerative protein misfolding and aggregation diseases such as Alzheimer’s Disease, Fragile X Syndrome, and Amyotrophic Lateral Sclerosis (ALS) (5-15). Understanding how G-quadruplexes exert power over aggregation could give clues to the roles of G-quadruplexes in these diseases.

Here, we explore the underlying biophysical and molecular determinants of the holdase activity of G-quadruplexes. We therefore systematically mutagenized known G-quadruplex sequences to determine important properties to holdase function, as has been previously performed with other chaperones (16). Structural investigations were then used to explore these properties in greater detail. We found that single point mutations of the G-quadruplexes can dramatically alter the holdase activity. Although the most important structural elements can vary depending on the G-quadruplex-containing sequence, structural dynamics or heterogeneity appears to be a common important feature of holdase activity.

## Materials and Methods

### Sourcing nucleic acids

All were ordered from Integrated DNA Technology using their standard desalting and purification procedures. For the heat aggregation plate reader assays, DNA was ordered lyophilized, and normalized to guaranteed molar weights by IDT. The DNA was then resuspended in 10 mM potassium phosphate buffer pH 7.5. For experiments using HEPES, DNA was transferred to the respective buffer with an Amicon spin column after 3 rounds of concentrating and washing with filtered de-ionized water and 3 rounds of concentrating and washing with the buffer it was being transferred into (eg 40 mM HEPES brought to pH 7.5 with KOH or LiOH).

### Annealing nucleic acids

Prior to every experiment, quadruplexes were annealed by heating to 95°C for two minutes and allowed to cool to 25 °C by a controlled temperature ramp of 1 °C/min with the (eppendorf ThermoMixer C). Samples were used immediately after annealing for all experiments.

### Thermal aggregation plate reader assays

For the initial thermal aggregation assays 550 nM citrate synthase from porcine heart (Sigma-Aldrich C3260-5KU) in a 1:2 protein:DNA strand concentration (1.1 µM DNA) in 10 mM potassium phosphate pH 7.5 or 40 mM HEPES brought to pH 7.5 with 6 M KOH or 2 M LiOH. Aggregation was measured via absorbance at 360 nm in a Biotek Powerwave XS2 multimode plate reader using black clear flat bottom half-area plates (Corning 3880), with shaking and measurements every 36 s for 90 minutes as previously described (3).

Briefly, for all assays, the plates were transferred from ice to a preheated 50°C plate reader, and the temperature was held constant throughout the entire experiment. The sequences were run in triplicate. Percent aggregation was calculated as a function of the maximum absorbance value recorded in the hour and a half divided by the maximum protein alone absorbance value. Error bars shown are standard error propagated from both the triplicate protein alone and triplicate experimental measurement. As a control, herring testes DNA (htDNA, Sigma D6898-1G) was also run on each plate to ensure consistency of data.

The relatively high melting point of the quadruplexes, all of them >60°C also indicates that the quadruplex would be intact and folded for our chaperone activity assay which takes place at 50°C.

### N-methylmesoporphyrin IX fluorescence

In a Corning 3880 plate, the emission spectra of 5 µM N-methylmesoporphyrin IX (NMM, Cayman Chemical) was measured using an excitation wavelength of 399 nm, and an emission of 610 nm in the presence or absence of 1 μM DNA in 10 mM potassium phosphate pH 7.5 or 40 mM HEPES brough to pH 7.5 with LiOH or KOH at a total volume of 100 µL per sample. Samples were run in triplicate at 25°C in a multimode plate reader (Tecan infinite M200). Reported values are the ratio of the emission at 610 nm for NMM + sequence divided by NMM emission in buffer alone.

### Circular dichroism and G4r-value

All CD spectra were measured using a Jasco J-1100 circular dichroism spectrophotometer. All samples were 22 µM DNA in 10 µM potassium phosphate pH 7.5 buffer.

For CD measurements at 25 °C, spectra were taken from 320 nm to 195 nm at 1 nm intervals using a 50 nm/min scanning speed, 8 second data integration time (D.I.T.), and the resulting spectra shown is the average of three accumulations.

To determine an individual sequence’s G4r-value, an equation from (17) was adapted where 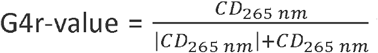 where CD _“### nm”_ represents the CD signal in mdeg at the given wavelength.

### QPCR Melting point analysis of LTR-III mutants

For determining the melting point of LTR-III and its mutants, a QPCR (QuantStudio 3) primer melting point analysis was used. To a room temperature 96 well, full skirt, clear PCR microplate (Axygen PCR-96-FS-C) a 50 µl solution of 1x SYBR Green (from 10,000x DMSO concentrate, MCE HY-K1004) with or without DNA in 10 mM potassium phosphate pH 7.5 were added. Once the samples were added, MicroAmp^™^ Optical Adhesive Film was used to seal the plate and ensure there was no evaporation from the sample. The samples were run in triplicate with measurements being made every 1° C from 25° to 95° C. The curves were analyzed via the QPCR software (QuantStudio^™^ Design & Analysis Software v1.5.1) and the first derivative of the melting curve was used to determine the melting point of a given sequence.

### Construction of expression vectors

pBAD33 vector was used to generate the expression vectors of G-quadruplex containing sequences, GroEL, and Empty (Fig 8A). pBAD33-GroEL and pBAD33-empty were obtained from previous study (3), and pBAD33mut-LTR-III WT and pBAD33mut-PARP-1 WT vectors were synthesized and cloned (Genscript) into pBAD33mut vector using the SacI and SpeI restriction enzyme sites. pBAD/HisD-TagRFP675 was a gift from Vladislav Verkhusha (Addgene plasmid # 44274; http://n2t.net/addgene:44274; RRID:Addgene_44274) (18).

### Protein expression, and Escherichia coli fluorescence assay

Each pBAD33 backbone-based vector and pBAD/HisD-TagRFP675 were co-transformed into the *E. coli* strain MC4100(DE3) by heat shock. Expression of protein and RNA was performed following the previous study (3). Briefly, each transformant was grown on a LB plate containing 0.2% L-Arabinose, ampicillin (200 μg/ml), and chloramphenicol (50 μg/ml) at 42°C overnight. To examine whether the expression is tightly controlled, LB plates containing the same amount of antibiotics were used for both +Empty and +GroEL samples, shown as NI. The next day, samples were prepared in 170 mM NaCl, and harvested by centrifugation at 10,000 *g* for 2 min at 4°C. Color of cell pellets was immediately compared after harvesting, and then cells were resuspended in the same buffer to 0.1 OD600 for further fluorescence assay. Fluorescence of TagRFP675 was measured by microplate reader (Infinite M200 Pro, Tecan) using a black 96-well plate (Corning Black NBS 3991) upon excitation at 598nm and emission at 675 nm.

### NMR sample preparation and data analysis

Oligonucleotides for NMR studies were procured in lyophilized form from Integrated DNA technologies (IDT USA), which were made with solid phase synthesizer and purified with desalting columns (standard desalting). Oligos were used without further purification. Concentration of DNA was estimated with UV absorbance at 260 nm, measured using Shimadzu UV-1800 UV-Vis spectrophotometer. G-quadruplex systems (LTR-III, LTR-III G10A, LTR-III TTTT) and hairpin duplexes (hp-G10A, hp-TTTT) were made to 9 mL (50 µM) concentraton and heated at 95 °C for 5 minutes, after addition of 1 mL of 0.2 M potassium phosphate buffer (pH 7.0), the soluton was mixed vigorously and heated again at 95 °C for 5 minutes. Samples were allowed to cool at room temperature for overnight. For making hp-stem duplex, a similar annealing protocol was used where the concentration of two complementary strands being 500 µM. Annealed samples were exchanged to NMR buffer (20 mM potassium phosphate, pH 7.0) using 3 kDa centrifugal filters (Amicon, Merck Inc.) spun at 4000 g and 4 °C to a final volume of 250 µL. NMR samples were made to 300 µL by adding 5% (v/v) D_2_O and 50 µM trimethylsilylpropanoic acid (TSP) as internal standard and NMR buffer. The final concentration of DNA samples were of 1-2.5 mM concentration. Samples were then transferred to 5 mm medium wall tubes (I.D 3.4 mm, Norell, Inc.).

NMR data was acquired with Bruker 700 MHz spectrometer equipped cryogenic triple ^1^H, ^13^C and ^15^N probe with z-gradients. All the data were acquired at 298K (and 310K for G4 systems) as decribed previously (4). ^1^H-^1^H NOESY (150/200 ms mixing time) (19), ^13^C-^1^H heteronuclear single quantum coherence (HSQC) (19) and ^15^N-^1^H heteronuclear multiple quantum coherence (HMQC) (20) spectra were acquired for each sample. Spectra were acquired with topspin 3.5.6 (Bruker Inc.), processed with NMRPipe (21) and analyzed using NMRFAM-SPARKY (22). Chemical shifts were referenced to TSP.

### Native PAGE Gels

All native PAGE gels were performed at 4°C with cold (4°C) 1X TBE running buffer (8.9 mM Tris base, 8.9 mM borate, with 2 mM EDTA) with 10 mM KCl or LiCl with Invitrogen 1.0 mm x 12 well 20% TBE-PAGE gels (EC63152BOX, Thermo Fisher Scientific). Gels were pre-run and run for 1 hour (unless otherwise stated) at 4°C once samples were added. Gels were first stained with 2 µM NMM (from a 10 mM stock in anhydrous DMSO stored at -20°C) in 1x TBE with 10 mM KCl or LiCl (to match gel running conditions) for 20 minutes and visualized with the SYBR green filter. Once visualized, the same gels were stained with 1x SYBR gold (from a 10,000x concentrate in DMSO) in 1x TBE with 10 mM KCl or LiCl and visualized with the SYBR gold filter. Gels were visualized on a BioRad ChemiDoc Imaging System.

Prior to running the gel, 50 µM nucleic acid stocks in appropriate counterion buffer (10 mM potassium phosphate pH 7.5 or 40 mM HEPES LiOH pH 7.5) were annealed using the above annealing protocol. Once annealed the samples were diluted to 5.6 µM in non-denaturing loading buffer (10% glycerol, 0.8 mM Tris base, and 9.6 mM glycine), and 1X TBE buffer with 10 mM KCl or LiCl. Immediately after dilution, 10 µL of sample was loaded into a lane and the gel was run and visualized according the procedure above.

For complete gel images see SI Figures 6-13.

## Results

### Single Point Mutations Dramatically Affect Chaperone Activity

To investigate why G-quadruplexes are such potent holdases, we began probing the activity of two mixed 3+1 G-quadruplexes, LTR-III and PARP-1. Our motivation for using these sequences is that they have solved tertiary structures and display multiple structural elements (23,24). Therefore, we could design mutants to disrupt particular structural features and test their importance in holdase activity. Although mutations were made to the G-quadruplexes to probe structural features; the G-quadruplex core was not altered, as we anticipated such mutations would completely destroy the known structure and obscure analysis. To evaluate the activity of the mutants we used a previously published thermal denaturation aggregation assay using the protein citrate synthase (CS) as a model client (3). To compare the holdase activity of the G-quadruplex mutants, we normalized the signal of each mutant to CS alone. The resulting ratio of the maximum aggregation level of a given mutant to that of CS alone is denoted as percent aggregation (%aggregation, Figure 2). Therefore, the most effective holdases will have a low percent aggregation.

**Figure 2.**
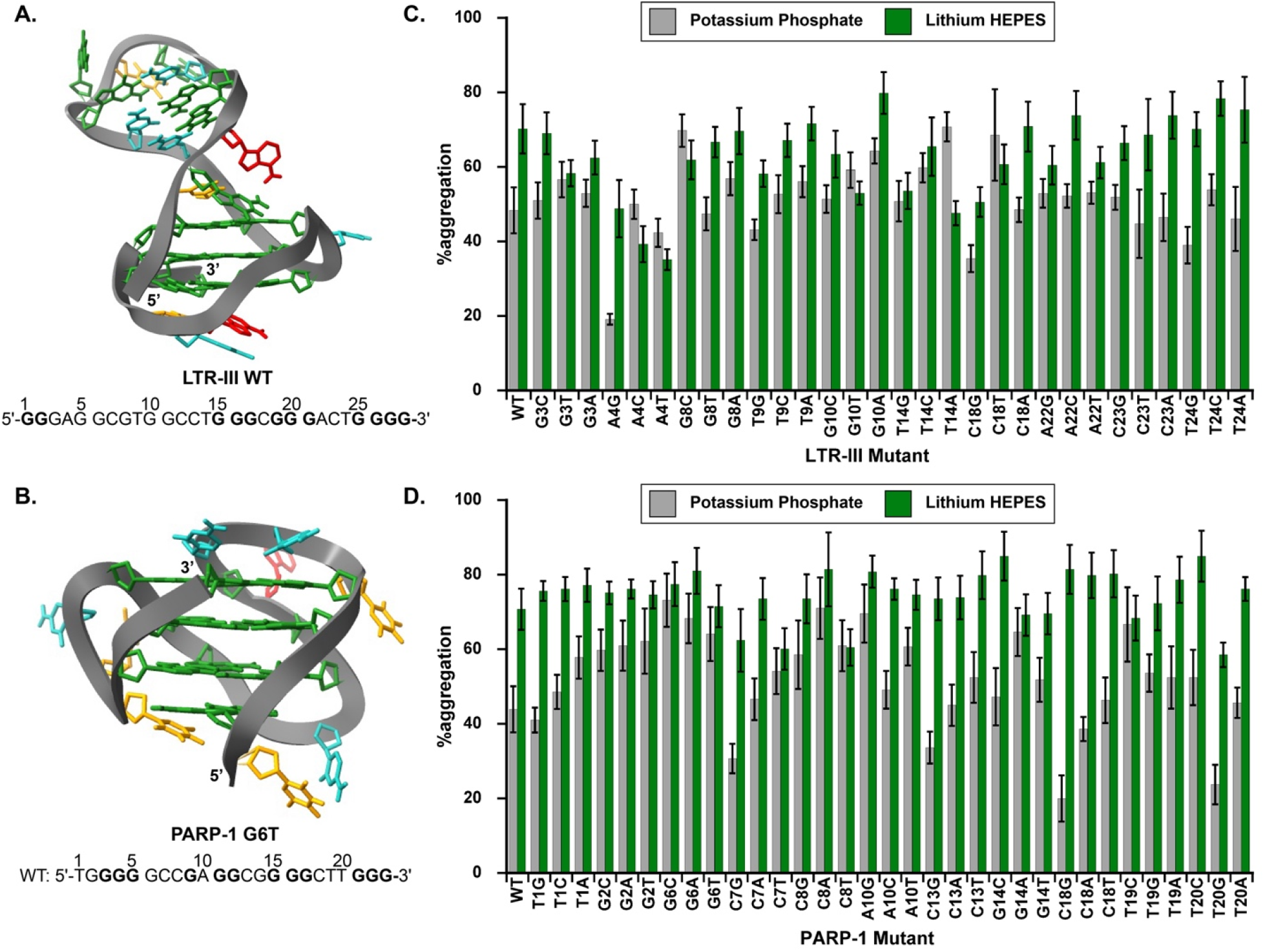
**A/B**. PDB-deposited NMR structures and wild type sequences for G-quadruplexes used here, bolded guanines are contained in the G-quadruplex core. **A**. PDB 6H1K. LTR-III WT structure and sequence. **B**. PDB 6AC7, an NMR structure of PARP-1 G6T and the sequence beneath is for WT PARP-1. **C./D**. Thermal denaturation aggregation assay results comparing %aggregation of 0.55 µM citrate synthase in the presence **C**. 1.1 µM LTR-III mutants and **D**. 0.275 µM PARP-1 mutants in either pH 7.5 10 mM potassium phosphate (grey) or pH 7.5 40 mM HEPES LiOH (green) buffers. Experiments were performed in triplicate and the error bars represent ± SE (see methods).

LTR-III WT has multiple distinguishing structural features. It is a 28-base DNA strand, containing a 12-guanine G-quadruplex core with a hairpin region extending from bases 5 to 13 (Figure 2A, PDB 6H1K). In 10 mM potassium phosphate buffer, about two-thirds of the LTR-III mutants we tested were within standard error of the LTR-III WT sequence (grey bars, error bars signify standard error Figure 2C), which had a 48% aggregation level. However, we did observe changes to holdase activity in both the positive and negative directions. Several of the mutations showed markedly worse holdase activity, including G8C, G10A, T14C, T14A, and C18T. Several of the mutations showed improved chaperone activity including A4G, C18G, and T24G compared to WT (Figure 2C). Together, the results found that simple single mutations could either increase or decrease holdase activity.

The other mixed 3+1 quadruplex, PARP-1, is a 23-base DNA sequence, (24) with a 12-guanine quadruplex core with a small capping region made up of G2, G14, and T20 (Figure 2B). Initially, PARP-1 and its mutants’ concentration used in the assay was identical to that of LTR-III’s (2:1 DNA:protein ratio), but PARP-1 WT and its mutants almost completely inhibited the aggregation of CS, with the resulting light scattering signal was essentially unchanged over the course of the 90-minute assay (data not shown). Although at this ratio, PARP-1 was highly effective, it made it difficult to compare how the point mutations affected chaperone activity. As a result, we screened lower PARP-1 concentrations and at a 0.5 to 1 DNA to CS ratio, we began to observe differential effects of the mutations (Figure 2D).

At this lower G-quadruplex concentration, clear activity differences between the mutants can be seen. PARP-1 WT has a %aggregation of ∼44%. Most of the mutations dramatically decreased the activity of the quadruplex, with almost 1/3^rd^ of the changes resulting in greater than 60% aggregation. Notably, any change made to the G2 or G6 positions resulted in markedly worse holdases. However, several highly potent holdases emerged, particularly C18G and T20G, both having %aggregation below 25% along with C7G and C13G both below 40% aggregation. (Figure 2D). Therefore, like LTR-III, simple point mutations cause dramatic changes in holdase activity in both directions. These experiments demonstrate the sensitivity of the holdase activity to changes in nucleic acid sequence or structure.

### Quadruplex Integral to Chaperone Activity

When looking at both G-quadruplexes, mutants that extended G-tracts in and adjacent to the G-quadruplex core were the strongest chaperones. For LTR-III these included A4G, C18G, and T24G. Similarly, all the most potent PARP-1 mutants: C7G, C13G, C18G, and T20G extended a G-tract adjacent to the G-quadruplex core. The inverse, disrupting G-tracts near the quadruplex core, were likely to be detrimental to chaperone activity, particularly for PARP-1. Any change made to G2, G6, or G14 of PARP-1 resulted in markedly worse activity compared to WT.

The above mutants suggest that G-quadruplex formation and stability is necessary for holdase function, particularly for PARP-1. To test this hypothesis, we took advantage of the differences in G-quadruplex stability in potassium or lithium buffers, as potassium stabilizes G-quadruplexes compared to lithium (25). To determine relative G-quadruplex formation, we used a fluorescent G-quadruplex binding dye, N-methyl mesoporphyrin IX (NMM) (26), and measured the fluorescent signal of the dye in both potassium and lithium buffers (Figure 3). An ideal substitute for our 10 mM potassium phosphate pH 7.5 buffer would be the equivalent lithium phosphate buffer; however, lithium phosphate is essentially insoluble at biological pH. Therefore, we substituted lithium phosphate buffer for 40 mM HEPES brought to pH 7.5 with lithium hydroxide (Lithium HEPES).

**Figure 3.**
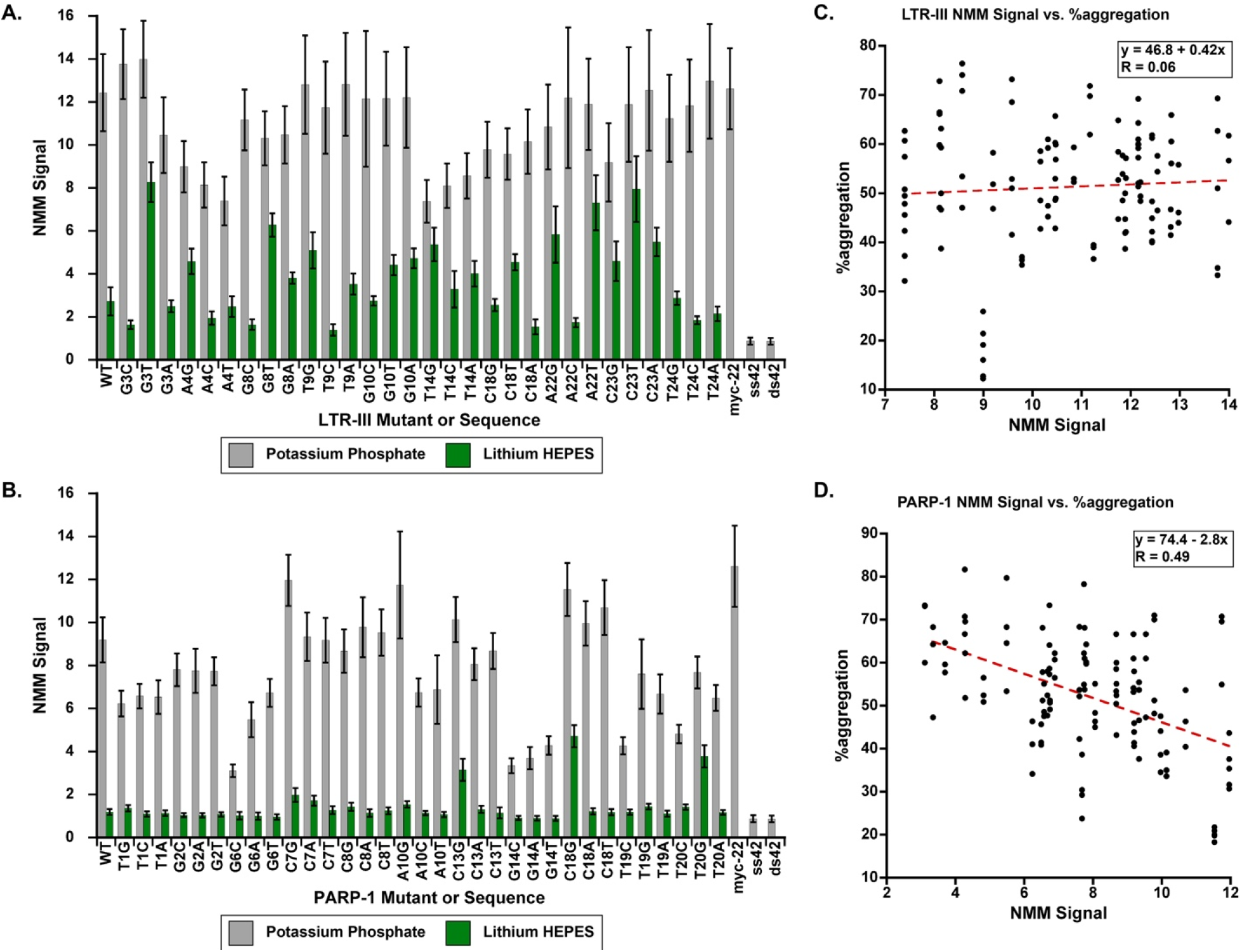
**A/B**. The ratio of the fluorescent signal of NMM + DNA was divided by the fluorescent signal of NMM alone to yield an NMM “turn-on” ratio we denote as NMM Signal (error bars are ± SE of triplicate data). **A**. NMM signal yielded by LTR-III mutants and **B**. PARP-1 mutants in potassium (grey) and lithium buffers (green). To verify the specificity of NMM G-quadruplex sequences a parallel topology forming G-quadruplex Myc22 served as a positive control and two negative controls were used: a single stranded DNA 20mer (ssDNA), and a duplexed version of the 20mer (dsDNA). **C/D**. Correlation plots between all %aggregation found for a given sequence (n≥3 technical replicates) and their NMM signal. **C**. LTR-III mutants were found to have no significant correlation (R = 0.06) and **D**. PARP-1 mutants yielded a negatively sloped curve with a moderate correlation (Pearson R value of 0.49), indicating stronger NMM signal is correlated with better holdase activity.

To directly compare between different experiments, the signal of NMM + G-quadruplex was divided by NMM alone to yield an NMM turn-on ratio (NMM Signal, Figure 3). As NMM doesn’t fluoresce in the absence of a G-quadruplex, its turn-on ratio will be 1 when there is no G-quadruplex present. To verify its specificity, we also tested two negative controls a single stranded DNA 20mer and a 20-base pair version, ssDNA and dsDNA respectively. Neither of these are G-quadruplex forming sequences and therefore NMM should not interact with either sequence. Our positive control was a 22-base parallel quadruplex forming sequence, Myc22 (27). NMM is thought of as primarily a parallel quadruplex-binding dye and therefore was expected to bind strongly to Myc22.

As can be seen in Figure 3A/B both LTR-III and PARP-1 mutants had dramatically reduced NMM signal in lithium compared to potassium. However, PARP-1 and its mutants are particularly sensitive to the cation substitution, as only six of the mutants were above the ssDNA and dsDNA controls in NMM signal. Notably, five of the six of these PARP-1 mutants substituted any base for guanine: C7G, A10G, C13G, C18G, and T20G (C7A being the sixth, Figure 3B). The NMM signal in potassium phosphate was noticeably higher for PARP-1 mutants, with most of the sequences having an NMM signal around or greater than 8 (Figure 3B). However, certain point mutations could greatly stabilize or destabilize the quadruplex. Changes to positions G6, G14, and mutants T19C, T20C all had markedly lower NMM signal than WT. However, several substitutions had the opposite effect and were noticeably higher than PARP-1 WT, including C7G and A10G along with C18 mutants (Figure 3B).

LTR-III WT and its mutants were still sensitive to the cation change, but the NMM signals in both lithium and potassium buffers were generally higher compared to PARP-1 mutants (Figure 3A/B). Analogous to the observations in PARP-1, LTR-III mutants that changed any base to cytosine appeared to be strongly destabilized by the presence of lithium: G3C, A4C, G8C, T9C, A22C, and T24C (Figure 3A). Each of these LTR-III mutants in lithium buffer had essentially the same signal as the ss/dsDNA negative controls. Interestingly, the location of these mutants does not seem follow any structural pattern, as the C-substitutions are found adjacent to or in the stem-loop or in loops connecting quadruplex planes. Although no structural pattern emerges, cytosine substitutions effectively increase G-C Watson-Crick base pairing competition within the structure which could inhibit G-quadruplex formation. In potassium, LTR-III WT and its mutants had a comparable signal to the Myc22 positive control. However, mutations to the two positions at the base of the stem-loop A4 or T14 had much lower signals relative to the other LTR-III sequences (Figure 3A.).

In addition to running the samples in potassium phosphate and Li HEPES, NMM signal was also measured in 40 mM HEPES brought to pH 7.5 with KOH (Potassium HEPES, SI Figure 2). This is a more direct comparison to determine if the change in buffer identity had any effect on the G-quadruplex stability. As can be seen in the SI Figure 2 the NMM Turn-on Ratio was unaffected by the identity of the buffer, with potassium HEPES and potassium phosphate having essentially the same ratio. Overall, the data shows that for PARP-1, the switch from potassium to lithium largely destroys the G-quadruplex structure. Whereas with LTR-III sequences, the G-quadruplex structure generally decreases, but is not reduced to baseline (Figure 3).

To determine whether the changes in the amount of G-quadruplex present with different cations affected activity, we tested the holdase activity of the mutants in the same Lithium HEPES for comparison (green vs. grey bars in Figure 2). Overall, the lithium buffer substantially decreased holdase activity. For PARP-I in Li HEPES, the lowest %aggregation across the sequences was 58% aggregation, ranging up to 85%. This is in sharp contrast to potassium where the average %aggregation increased from 52% in potassium buffer to 74% in lithium (Figure 2D). The sequences that were most strongly affected were the most potent holdases in potassium: C7G, C13G, C18G, and T20G; all of which had %aggregations 2 to 4x higher in lithium compared to potassium buffer.

For LTR-III WT and its mutants, the switch to lithium generally decreased holdase activity, but the effects were considerably less dramatic compared to PARP-1 (Figure 2). The mutants most strongly affected by the switch to lithium were LTR-III A4G, T9G, C18G, A22C, and position T24. Several of these were the most powerful holdases in potassium, specifically A4G, C18G, and T24G. Overall, the substitution of potassium for lithium was highly detrimental to the holdase activity of the quadruplex forming sequences, particularly for PARP-1, indicating the importance of the G-quadruplex structure for holdase activity.

To further validate the importance of G-quadruplex structure to holdase activity, we plotted NMM Signal for each of the mutants against their %aggregation across all their replicates (n ≥ 3 technical replicates). To determine if a correlation existed between NMM signal and holdase activity, we subjected the data to a linear regression to determine the Pearson R value (Figure 3 C/D). For LTR-III mutants, there was zero correlation with a Pearson’s R of 0.06 (Figure 3C). However, for PARP-1 a moderate correlation between NMM Signal and %aggregation emerged, with a Pearson R of 0.49. As shown in Figure 4b, higher NMM values correlate with lower %aggregation values, indicating that stronger NMM binding is associated with stronger holdase activity. Although intriguing, NMM does not necessarily bind all G-quadruplexes with the same affinity, with a preference for parallel G-quadruplexes. We therefore continued to explore the dependence on specific G-quadruplex topology using circular dichroism (CD) spectroscopy.

**Figure 4.**
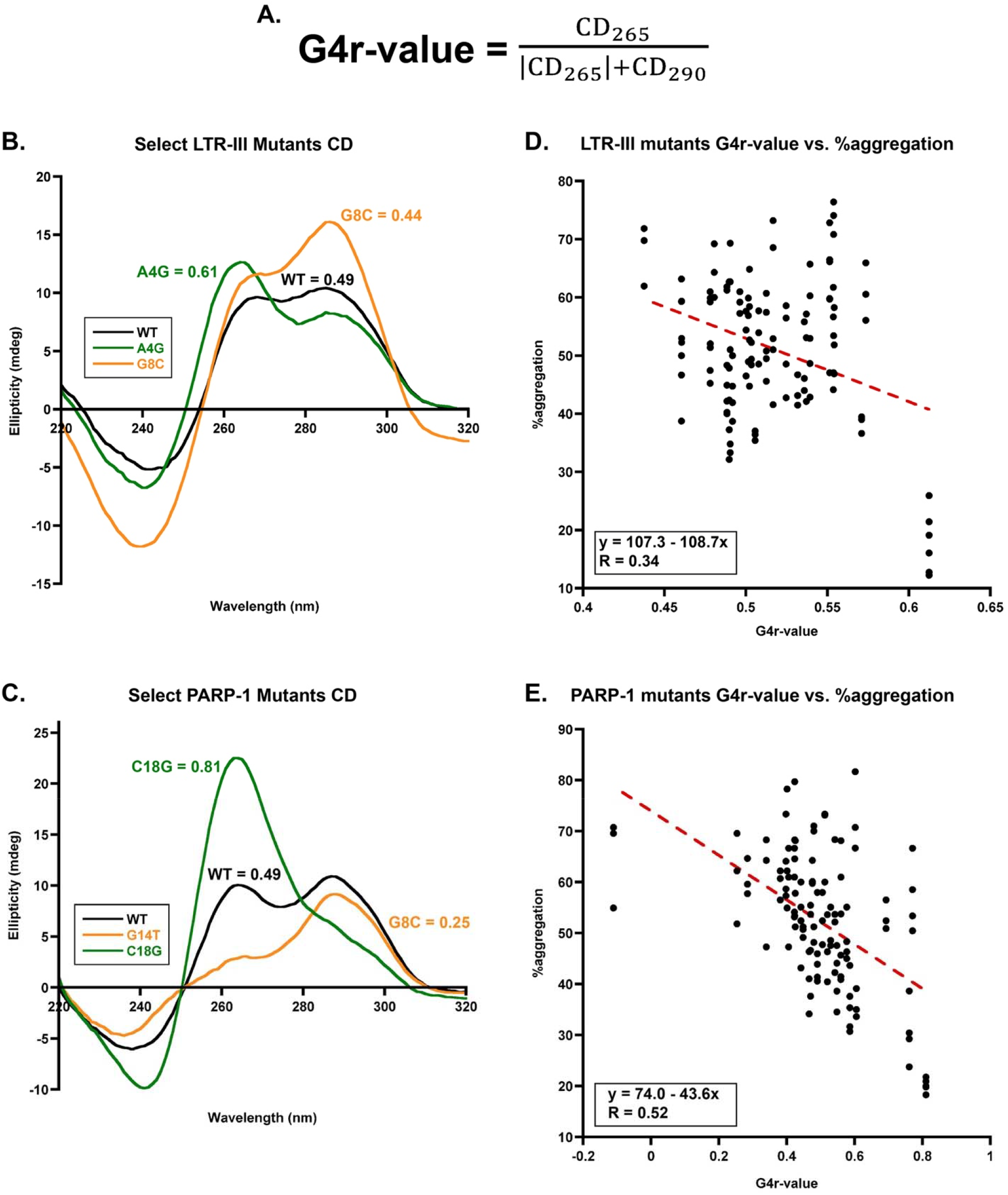
**A**. The G4r-value is a means to measure the relative topology of a given G-quadruplex forming sequence via CD. Subscript numbers indicate CD signal in mdeg at those given wavelengths in nm. **B/C**. CD spectra of select **B**. LTR-III mutants and **C**. PARP-1 mutants with G4r-value shown. **D./E**. Correlation plots between all %aggregation found for a given sequence (n≥3 technical replicates) and their G4r-value. For both LTR-III mutants **D**. and PARP-1 mutants **E**. a correlation emerged where G-quadruplexes with more parallel topology (G4r-value closer to 1) correlated with stronger holdase activity. The Pearson R value of 0.34 for LTR-III mutants indicates a weak correlation while PARP-1 yielded a moderate correlation with a Pearson R of 0.52. Note the position of the y-axis in **E**., one of the PARP-1 mutants, A10G caused the mixed 3+1 G-quadruplex to convert to a fully anti-parallel topology resulting in a negative G4r-value (see SI Figure 4 for full spectra).

### Dependence of holdase activity on G-quadruplex topology

A lingering question from the chaperone activity assay was if the point mutations were making large changes to the quadruplexes’ topology. To test this possibility, we performed CD spectroscopy on all the sequences for both G-quadruplexes. As can be seen in Figure 4A/C (SI Figure 3 and SI Spreadsheet) the structure of many of the G-quadruplexes is relatively similar to the WT and all but one of the sequences contained the mixed 3+1 G-quadruplex characteristic maxima at 260 and 295 nm. Interestingly, one of the poorer performing PARP-1 sequences A10G induced a topology change from mixed 3+1 to anti-parallel (SI Figure 4). This is consistent with our prior data indicating that anti-parallel quadruplexes are generally poorer holdases compared to their mixed 3+1 counterparts (3). In general, these CD spectra indicate the mutations are not disrupting the G-quadruplex core and that the differences observed in holdase activity are directly comparable between mutants with relatively similar structural properties.

To further determine the relationship between G-quadruplex topology and holdase activity, we used the 290 and 265 nm CD signals to quantify the topology of the G-quadruplexes (Figure 4B/C). We used a previously published method that quantitates the amount of parallel and anti-parallel character of a G-quadruplex forming sequence using these signals as a ratio, referred to originally as an r-value, here as a “G4r-value” (17). To determine the relative topology of the quadruplexes used here, a G4r-value of 0.5 is treated as the ideal mixed 3+1 quadruplex with equal peaks at 290 and 265 nm. A G4r-value closer to 1 indicates the quadruplex is more parallel, and closer to zero indicates more anti-parallel. If a G4r-values is negative (< 0) the G-quadruplex has an anti-parallel topology and if a G4r-value = 1 it is parallel (17).

We plotted the %aggregation against the G4r-value and used Pearson R to quantify if there was any correlation. For LTR-III and its mutants, a weak correlation between %aggregation and G4r-value emerged with a Pearson R of 0.34 (Figure 4D). This indicates that a G-quadruplex with more parallel topology character was weakly correlated with lower %aggregation and stronger holdase activity. However, the median G4r-value for LTR-III was 0.51 ranging from 0.44 to 0.61 indicating they had relatively little parallel or anti-parallel character across the mutants and were primarily mixed. For PARP-1, the same trend emerged where more parallel character correlates with stronger holdase activity with a moderate Pearson R of 0.52 (Figure 4E). PARP-1 mutants also had a much larger range of data, from completely anti-parallel (negative data points) to G4r-values as high as 0.8, which is almost completely parallel. Both the weak correlation of LTR-III and moderate correlation of PARP-1 agrees with prior data indicating that generally anti-parallel quadruplexes are poor holdases and parallel quadruplexes generally make stronger holdases.

### Hairpin dynamics affect holdase activity of LTR-III

To isolate the contributions of the quadruplex core in LTR-III WT, we created a variant that exchanged the stem loop for a 4-thymine diagonal loop across quadruplex plane-3. (LTR-III TTTT mutant Figure 5). Interestingly, this core alone recapitulated similar holdase activity to LTR-III WT with a %aggregation of 50% showing that the G-quadruplex core is essential to holdase activity. NMR spectroscopy confirmed the formation of a G-quadruplex core similar to LTR-III WT in this variant (Figure 5).

**Figure 5.**
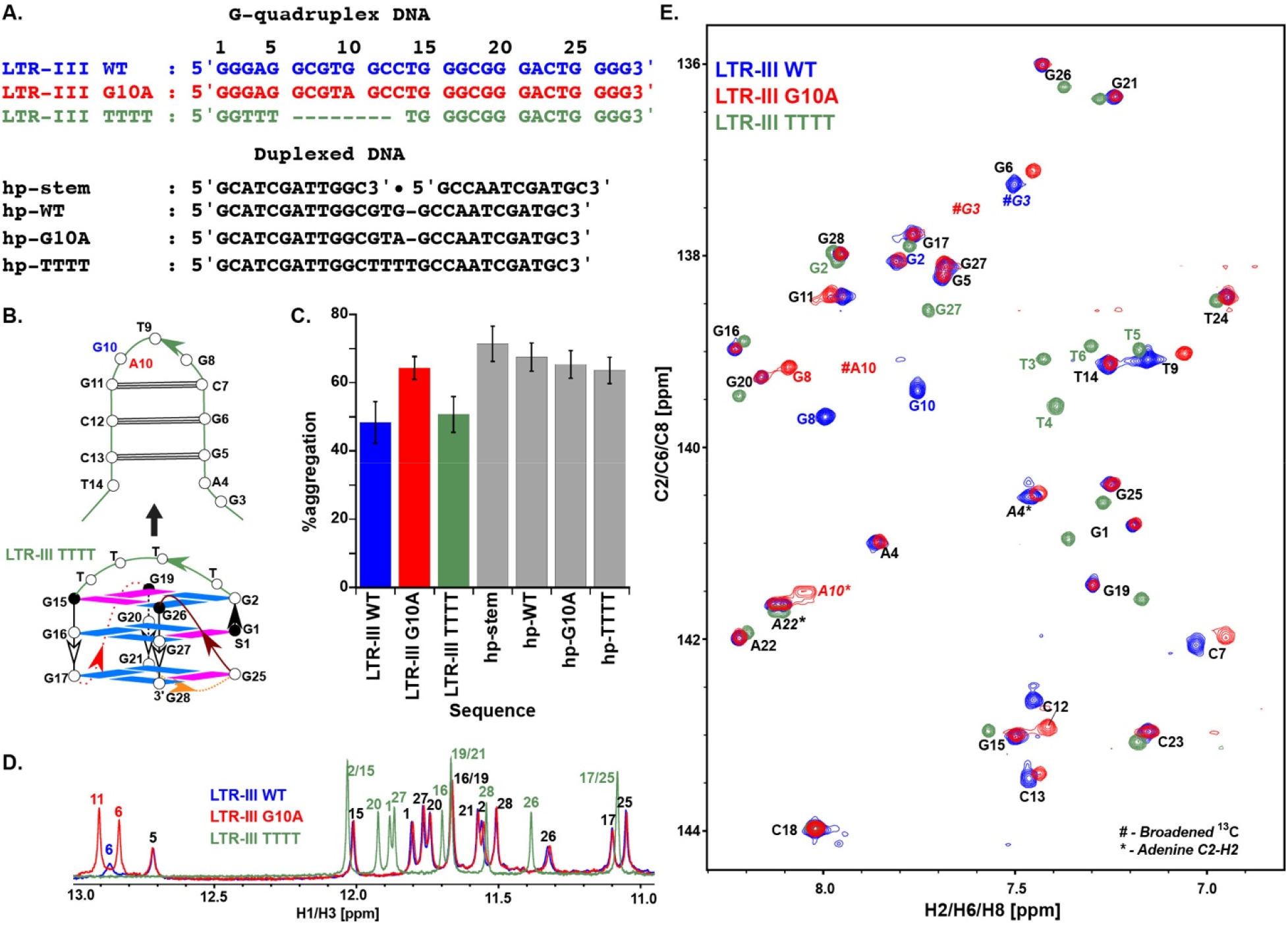
**A**. G-quadruplex and duplexed DNA sequences used for thermal denaturation aggregation assay and NMR studies. Regions are highlighted to show sequence similarity between sequences and LTR-III WT. **B**. Schematic for duplex constructs and LTR-III WT-based G-quadruplex constructs. **C**. Thermal denaturation aggregation assay comparing %aggregation vs DNA construct (error bars ± SE). **D**. Overlay of ^1^H-1D imino resonances LTR-III WT (blue), LTR-III G10A (red) and LTR-III TTTT (green). LTR-III TTTT and G10A displayed 12 peaks in their imino ^1^H spectrum, indicating formation of the G-quadruplex core, with NOESY data confirming stable diagonal loop formation. **E**. Overlay of ^13^C-^1^H region (aromatic) resonances of LTR-III WT (blue), LTR-III G10A (red) and LTR-III TTTT (green). For LTR-III TTTT and G10A the ^13^C/^1^H shifts confirm the formation of hybrid (3+1) conformation similar to that of LTR-III WT. **D and E**. Closely matched resonances are marked in black and all other resonances in respective colors and numbering indicates positioning according to LTR-III WT sequence. In **E**. # indicates broadened resonances in ^13^C dimension and * indicates ^13^C aliased resonances of Adenine C2-H2.

Despite the importance of the G-quadruplex, in LTR-III WT and its mutants the hairpin appears to be an important contributor to its chaperone activity. Several of the least powerful LTR-III holdase mutants G8C, G10T/A, and T14A along with one of the strongest holdases, A4G, were immediately adjacent to the hairpin or a part of it. Therefore, we hypothesized that the stem-loop conformational preferences contribute to the holdase activity.

LTR-III G10A had previously been suggested to have different hairpin dynamics than WT (23), but had a similar CD spectrum suggesting very similar structural characteristics to WT (SI Figure 3), and lower holdase activity than WT LTR-III (Figure 2C). To determine if the loop or the stem of the hairpin were contributing to the chaperone activity of LTR-III, we created a series of duplexed hairpin variants without the G-quadruplex core: a hairpin version of the WT stem-loop (hp-WT), a version of the WT stem without a loop (hp-stem), a hairpin with an all thymine loop (hp-TTTT), and a hairpin of mutant G10A (hp-G10A) (Figure 5A/B). Duplex formation was confirmed by NMR (SI Figure 5). Despite all of these variations, none of the duplexed variants displayed substantially different activity from each other, suggesting that the loop on its own does not substantially govern holdase activity (Figure 5C). Therefore, the effect of the G10A mutation on holdase activity requires the native structural context and it is possible this is also true for other hairpin variants.

To examine the role of the loop dynamics, we performed conventional 1D and 2D NMR experiments with WT LTR-III and LTR-III stem-loop mutant G10A. We hypothesized that the dynamics of the loop could influence the overall structural context of LTR-III and effect holdase activity.

Excellent overlap of imino ^1^H resonances (Figure 5D) were observed between LTR-III WT and LTR-III G10A (Figure 5A and 5C), confirming similar overall structure (23). The 10-12 ppm range unequivocally confirmed that the G10A mutation maintained the intact G-quadruplex core, as reported earlier (23). Furthermore, ^13^C/^1^H chemical shifts for LTR-III WT validated the hybrid (3+1) topology (with the characteristic v-loop) and LTR-III G10A maintained the same architecture (Figure 5E, adjudged from 8:4 guanosine nucleotide in *anti:syn* ratio) (4,23).

Closer inspection of the NMR spectra of LTR-III WT and the G10A mutant shows noticeable differences in the non-tetrad regions of the spectrum. Narrow imino ^1^H resonances observed for G6 and G11 in LTR-III G10A indicates a more stable Watson-Crick paired helix in the stem than in the case of LTR-III WT (broad resonance of G6 observed, Figure 5D). This observation suggests that in WT LTR-III, G10 competes with G11 to form a pair with C7. This base-paring competition is removed by the G10A mutation. This observation is consistent with the broadened ^13^C-^1^H resonances (in the range of 298-310K) in the 2D NMR spectrum (Figure 5E) for A10, suggesting increased A10 dynamics and flexibility compared to G10.

Overall, it appears this substitution G10A decreases global dynamics of the stem-loop in spite of the individual A10’s increased local dynamics compared to WT. Prior studies have shown greater chaperone dynamics is associated with stronger activity (16). A similar phenomenon could be occurring here as a result of the shifting base-pairing of the hairpin in LTR-III WT, leading to overall increased stem-loop dynamics that can enhance holdase activity.

In prior NMR work by the Phan group, it was shown that the PARP-1 WT sequence adopts a range of G4 conformations, while a single mutation, G6T, results in the formation of a stable hybrid (3+1) G4 topology without any flexible loops being present (4,24). In our study the PARP-1, the G6T mutant resulted in a significant reduction in holdase activity compared to PARP-1 WT (Figure 2D). In combination with our observations on LTR-III WT and G10A, we propose that the combination of a G-quadruplex scaffold in combination with more conformationally dynamic regions drive holdase activity for G-rich sequences.

### LTR-III Mutant Stability and Holdase Activity

The NMR experiments demonstrated that changes in the structural dynamics of LTR-III changed its holdase activity. To approximate changes in dynamics from other mutants, we measured melting temperatures for LTR-III mutants. This approach is analogous to that previously used to analyze the role of structural dynamics in the chaperone Spy (16). In this previous study, there was a negative correlation between melting temperature and activity. Although melting temperature and dynamics are not perfectly correlated, previous studies of nucleic acid dynamics suggest that nucleic acid motions and non-native conformations typically increase with increased temperature (28,29).

Overall, single point mutations were able to yield a melting point as much as 3 °C lower (C18G) than WT LTR-III and as much as 11 °C higher (Figure 6a and SI Table). The G10A mutant, which as discussed above had reduced hairpin conformational dynamics and was one of the worst performing holdases and had among the highest melting points.

**Figure 6.**
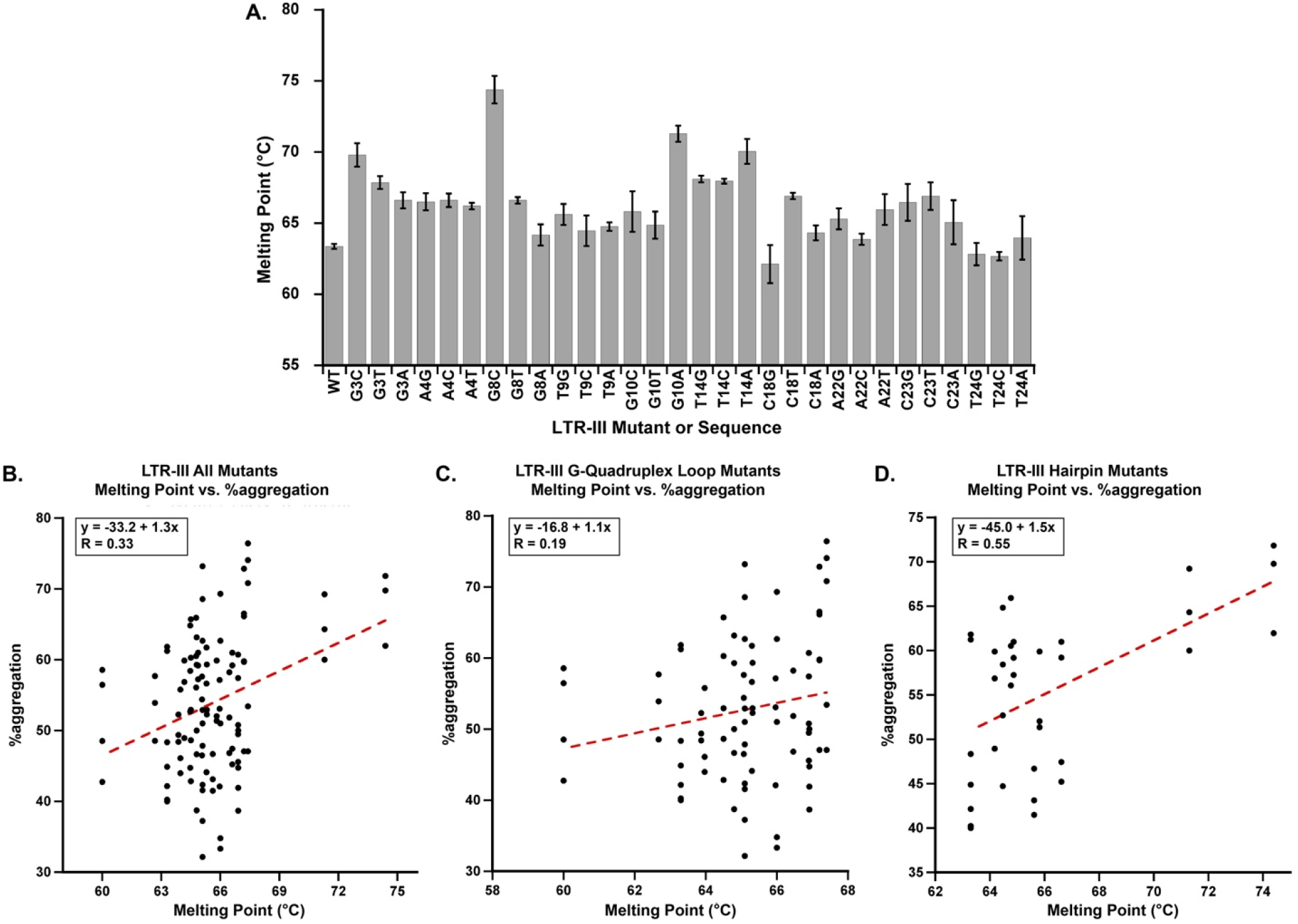
**A**. The melting points of LTR-III mutants determined from a qPCR primer melting point assay using SYBR green fluorescence to monitor unfolding (error bars are ± SE). **B, C, and D**. Correlation plots between all %aggregation found for a given LTR-III mutant (n≥3 technical replicates) and their melting point. All plots include LTR-III WT and exclude A4G, C18G, and T24G as their heterogeneity made it difficult to determine the dominant species contributing to the melting temperature (see Figure 7). **B**. LTR-III mutations made to both the stem-loop and to bases at the periphery of the G-quadruplex core. A Pearson R value of 0.33 indicates there is a weak correlation between higher melting points and reduced holdase activity across all mutants. **C**. LTR-III mutations made only to the periphery of the G-quadruplex core. A Pearson R value of 0.19 indicates there is a very weak correlation between the melting point of these mutations and holdase activity **D**. The correlation for LTR-III mutations made to only to the stem-loop. The linear regression yielded a positively sloped correlation between higher melting points and decreased holdase activity.

To see if dynamics play a part in holdase activity, we plotted melting temperature and %aggregation for all LTR-III mutants (minus oligomerically heterogeneous mutants covered in the next section). This found a weak correlation (Pearson R = 0.33) in which increased melting point was correlated to decreased holdase activity (Figure 6B). However, it was unclear if mutations made to the hairpin or to the peripheral loops on the G-quadruplex itself could be more correlated to activity, so we plotted them separately. When we plotted the melting temperature vs. %aggregation for the mutations made to the periphery of the quadruplex core of LTR-III, we saw essentially only a very weak correlation between activity and stability (Pearson R of 0.19, Figure 6C). On the other hand, the hairpin loop mutations displayed a moderate positive correlation between the melting temperature and %aggregation with a Pearson R of 0.55 (Figure 6D). On its own, this comparison suggests mutations in the hairpin that stabilize the structure are correlated to less powerful holdases, consistent with the NMR analysis.

### Heterogeneity and chaperone activity

Despite observing changes in holdase activity that depend on topology and stability, other factors could still influence holdase activity. To examine whether higher order structure could also play a role, we ran native gels to determine if some of the mutants produced larger oligomeric structures. In prior literature, PARP-1 WT was shown to be relatively unstable, readily forming larger oligomers while LTR-III was shown to be largely monomer (also indicated by the NMR work here). The native gels were visualized with a G-quadruplex-specific and a non-specific nucleic acid stain (NMM and SYBR Gold, respectively).

When observing the WT of each G-quadruplex forming sequence, there is large discrepancy in the number of bands present for each. Each sequence has a large band corresponding to monomer, but PARP-1 WT has at least 5 bands and a large degree of smearing compared to LTR-III WT, which has only 1 additional faint band above it. This pattern holds true for all the PARP-1 sequences (select gels in Figure 7C), where there is generally a greater degree of smearing and number of bands compared to LTR-III WT and mutants.

**Figure 7.**
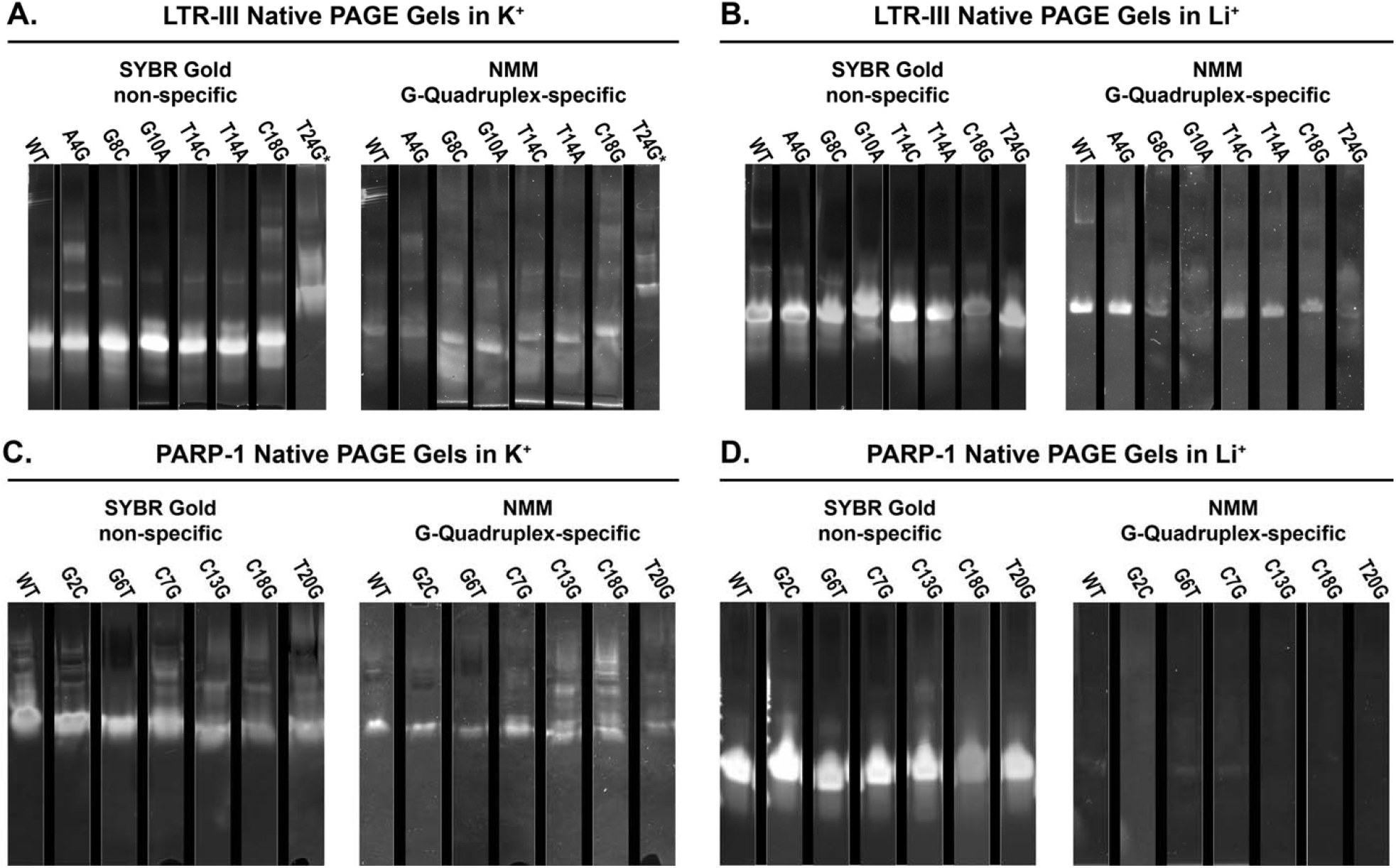
Non-denaturing native PAGE gels of select **A/B**. LTR-III mutants or **C/D**. PARP-1 mutants in potassium **A/C**. or lithium containing TBE buffer **B/D** stained with SYBR gold (left), a non-specific DNA dye or NMM a G-quadruplex specific dye (right). Going from potassium to lithium containing buffers reduces the overall number of NMM positive bands for heterogenous LTR-III mutants, this suggests less stable larger G-quadruplexes are lost in the absence of potassium. (**A/B**. See A4G and C18G) In contrast, almost all NMM-positive bands for PARP-1 mutants are absent in the presence of lithium, indicating the G-quadruplexes are not detectable in the absence of potassium (**C/D.**). T24G* indicates the gel was ran for 1 hour compared to 1.5 hours for the other LTR-III potassium containing gels, leading to a higher monomer band.

When we look more loosely at the gels for the PARP-1 mutants, the number of bands and degree of smearing can be loosely correlated to its holdase activity. For example, the most poorly performing PARP-1 mutants at positions G2 and G6 all had markedly reduced number of bands and smearing compared to WT. In contrast, the strongest holdases C7G, C13G, C18G, and T20G all had bands above the monomer band had with a high degree of smearing. The same pattern applies to LTR-III, as the most potent mutants, A4G, C18G, and T24G all have more bands than WT. These higher molecular weight species for both G-quadruplexes are NMM-positive. Together, this data suggests that larger intermolecular G-quadruplexes are likely responsible for their enhanced activity (Figure 7).

In the thermal denaturation aggregation activity assay described above (Figure 2), the holdase activity of PARP-1 WT and its mutants was negatively impacted by the switch from potassium to lithium counterion. The change was less detrimental for LTR-III and its mutants overall, however, several LTR-III mutants that contained multiple bands on the native gels were the most negatively impacted by the switch to lithium (LTR-III A4G and C18G, Figure 7A). As we had already determined that lithium destabilized PARP-I but only partially destabilized LTR-III structure via NMM binding (Figure 3), we sought to visualize how the metal change would affect the G-quadruplex oligomerization by running the same native gel experiments in lithium buffer.

For LTR-III, the G-quadruplexes can still readily form in lithium as they are NMM-positive, but the change in metal greatly reduced the number of oligomeric G-quadruplexes present. In lithium, the samples are primarily monomer G-quadruplexes, even for mutants that previously had multiple bands in potassium, such as A4G, C18G, and T24G (Figure 7B). Notably, these were the strongest holdases, but in lithium buffer, they specifically lost their powerful activity to a greater extent than other mutants. Together, these data indicate that the higher activity of these most powerful LTR-III mutants stemmed primarily from their higher order structure, similar to PARP-I.

### LTR-III Improves TagRFP675 folding in *E. coli*

In prior work we used a relatively unstable fluorescent protein, TagRFP675 as an in-cell folding sensor. Like other fluorescent proteins, TagRFP675 does not fluoresce at its excitation maximum unless it is in its native state, and in the absence of co-expressing chaperones we have shown that the protein has very little fluorescent yield (Figure 8B). To observe if the G-quadruplexes tested here had any effect on the folding and fluorescent yield of TagRFP675 in *E. coli* we co-expressed TagRFP675 with wild type PARP-1 and LTR-III.

**Figure 8.**
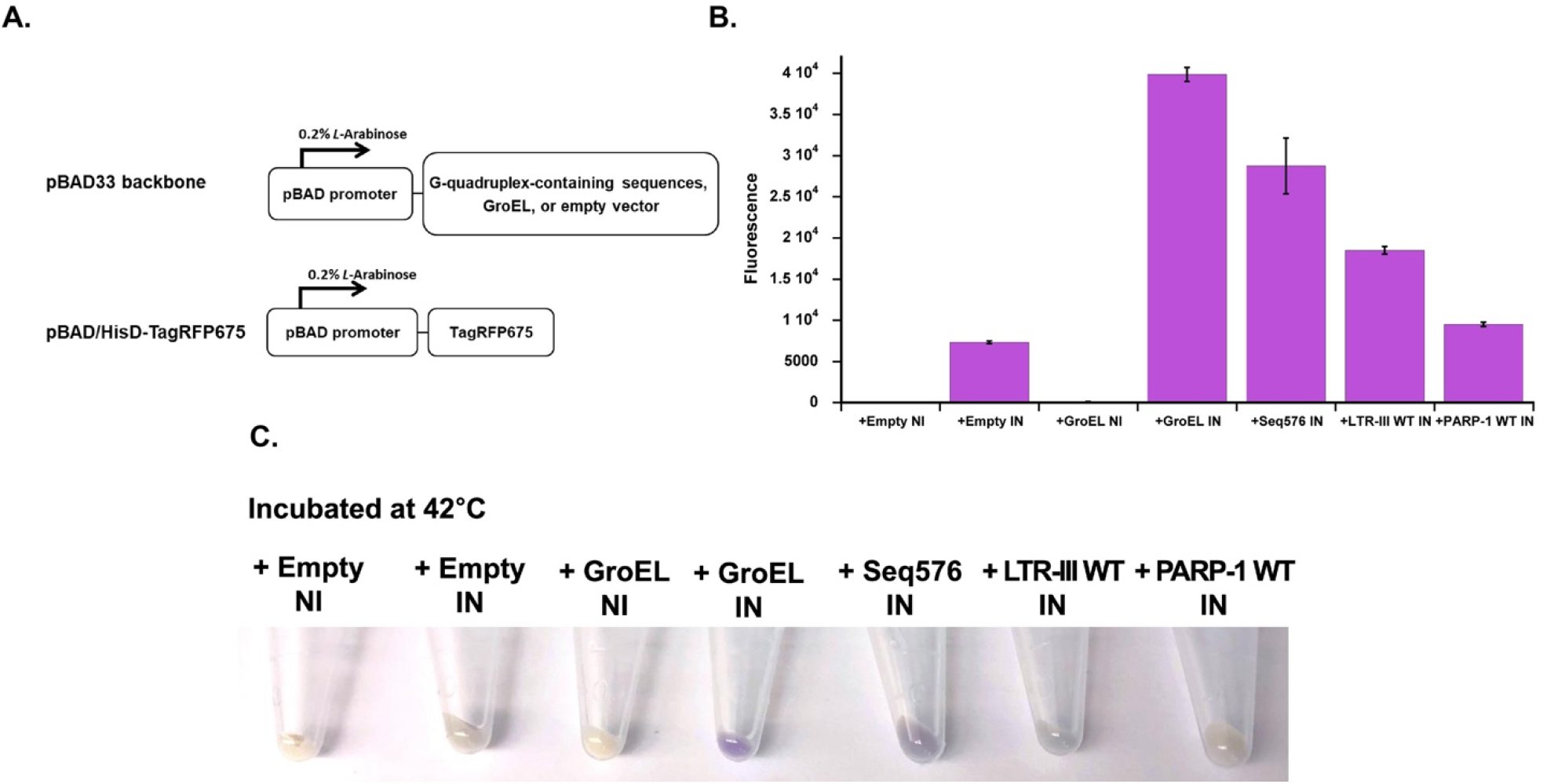
**A**. A simplified model of expression vectors used here. The expression of an empty vector, GroEL, and G-quadruplex forming sequences: PARP-1 WT, LTR-III WT, and the previously studied G-quadruplex Seq576 are under the control of an *L*-arabinose (0.2%) inducible pBAD promoter. **B**. Cellular fluorescence assay of TagRFP675 with negative controls (+Empty NI and IN, and GroEL NI), positive controls (+GroEL IN and +Seq576 IN), and G-quadruplex-forming sequences LTR-III WT and PARP-1 WT. Protein expression was induced at 42°C. The experiment shown here were technical triplicates (n = 3) and performed more than 2 times. **C**. Harvested cells showing relative color of *E. coli* pellets, more purple pellets correlate to stronger fluorescent signal. NI and IN indicate non-induced and induced, respectively.

We see very little fluorescence when TagRFP675 is induced without a co-expressed chaperone. However, when we induce TagRFP675 with chaperones we see a large difference in the amount of fluorescence compared to the negative controls. Consistent with prior results, co-expressing GroEL creates large amounts of folded protein as does a previously studied parallel G-quadruplex forming sequence, Seq576 that is reflected as a higher fluorescence signal. When we focus on LTR-III WT and PARP-1 WT, we see that LTR-III has a much higher signal than PARP-1, which displays little fluorescence (Figure 8B). These results are qualitatively supported by simply pelleting out the *E. coli* from these various conditions (Figure 8C). There is clearly a much darker purple color associated with the strongest chaperones: GroEL, Seq576, and LTR-III compared to the predominantly paler yellow color of the negative controls and PARP-1.

This result is interesting in that from everything we have shown so far, PARP-1 and its mutants have been the more potent chaperone than LTR-III *in vitro*. However, there are several factors that likely lead to it being a less potent chaperone in *E. coli*. From the NMM-fluorescence binding assays as well as the NMM-positive gels (Figure 3 and 7), we can see that LTR-III WT has a much stronger signal than PARP-I WT, suggesting perhaps it forms quadruplex structure more stably especially under non-ideal conditions. Similarly, we also have shown that in the absence of potassium that NMM is unable to bind PARP-I, suggesting the G-quadruplex is essentially destroyed. It therefore seems likely that in the complex environment of the cell, perhaps PARP-I WT is not forming as stable a G-quadruplex as LTR-III WT and therefore unable to provide significant chaperoning power.

## Discussion

In summary, here we determined several important aspects of G-quadruplexes that drive their ability to prevent protein aggregation. We first showed that the holdase activity of G-quadruplexes could be altered positively and negatively by single mutations. We then showed that the G-quadruplex core was essential for activity via metal substitutions. From CD spectroscopy and NMM binding experiments, we showed that the topology contributed greatly to the activity. Through a combination of NMR and melting experiments we uncovered the contribution of dynamics to the holdase activity. Finally, we found that in many cases that the oligomer state of the G-quadruplexes appears to also influence activity. These may still not be all of the important factors that can contribute to activity, but all of these appear to be major players.

The dependence of the holdase activity on structural topology was previously hinted at by examining a small number of parallel, anti-parallel, and mixed G-quadruplexes (3), but it was unclear whether other complicating factors could have biased the small sample size and large changes in structure. The more thorough examination here strongly indicates a topology dependence for holdase activity. However, why the structural topology drives holdase activity is less clear. There could be several underlying driving forces, including different oligomeric structure, change in binding stoichiometry, among others.

The finding that oligomerization was important to activity was not surprising, as we had previously seen the formation of protein:nucleic acid oligomers as a regular phenomenon among nucleic acids acting as chaperones (30), and G-quadruplexes in particular (3). We had previously found that this oligomerization could at times act as kinetic traps to prevent further aggregation, but sometimes could accelerate overall protein aggregation. Given the nucleic acid itself oligomerizes could provide a mechanism for co-oligomerization with protein clients.

As chaperones need to interact with mobile clients, dynamics are a common feature of many chaperones (31). Although some of these dynamics are large scale motions driven by ATP hydrolysis, the ATP-independent chaperone Spy also has dynamics as an important component of its mechanism (16,32). The oligomerization of G-quadruplexes is also reminiscent of the mechanism of small heat shock proteins (33). It is possible that the oligomerization and dynamics are both feeding from a similar entropic driving force to maximize binding sites.

While the role of G-quadruplexes protein aggregation diseases is still largely unclear, links now show them playing roles in diseases such as Fragile X syndrome, ALS, and Alzheimer’s disease. The structural features identified here could help in determining the role of G-quadruplexes implicated in these diseases.

## Data Availability

All data except NMR spectra available in Supplementary Table and Supplementary Information. For raw NMR data, please contact the authors.

