## Supplementary Data for "Why are G-quadruplexes good at preventing protein aggregation?"

**Supplemental Information**

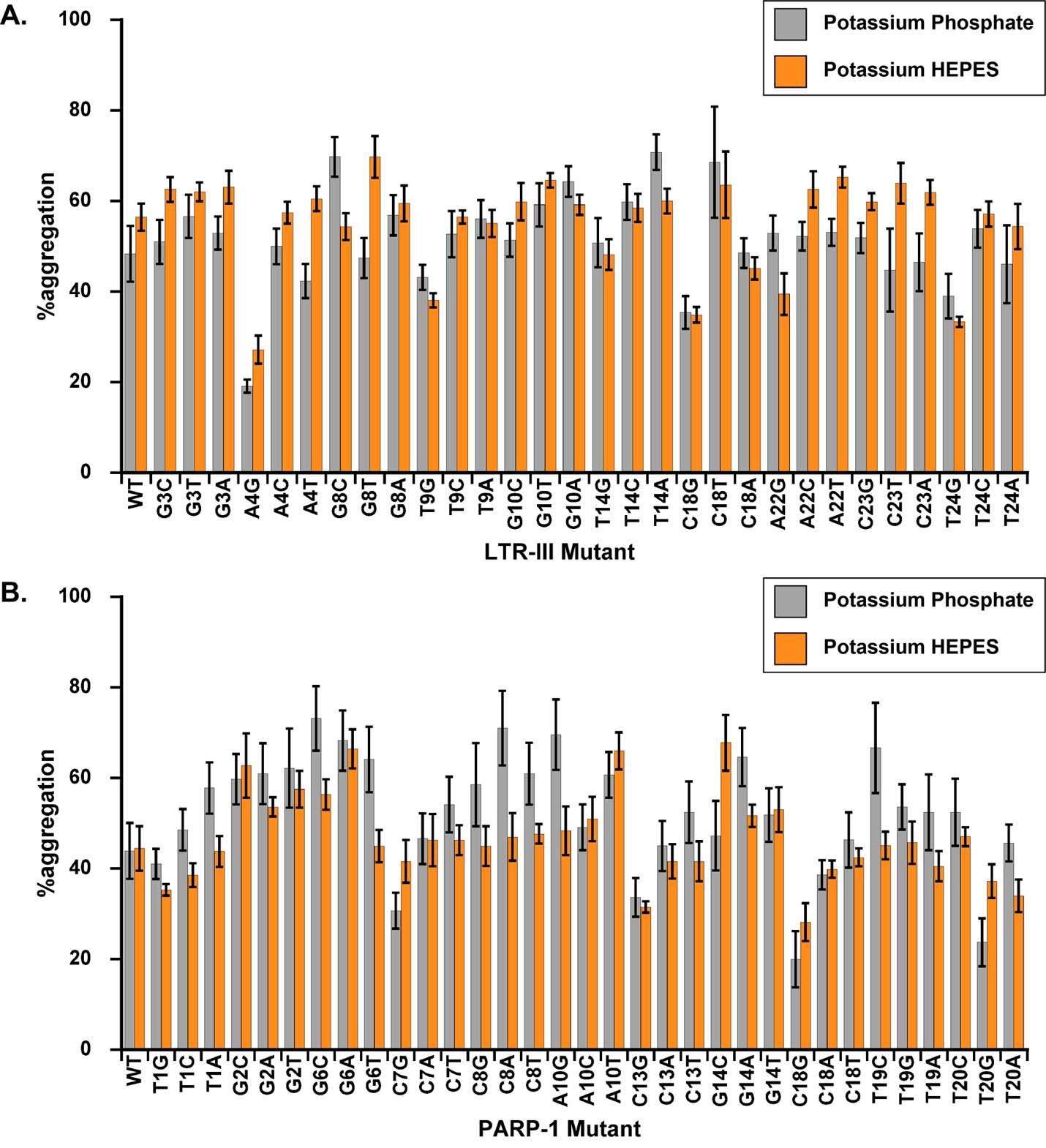

**SI Figure 1.** %Aggregation results from thermal denaturation aggregation assay comparing **A.** LTR-III and its mutants or **B.** PARP-1 and its mutants in 40 mM HEPES KOH buffer pH 7.5 to 10 mM potassium phosphate buffer pH 7.5. The relative %aggregation for individual mutations and comparisons between mutations is consistent between the potassium containing buffers.

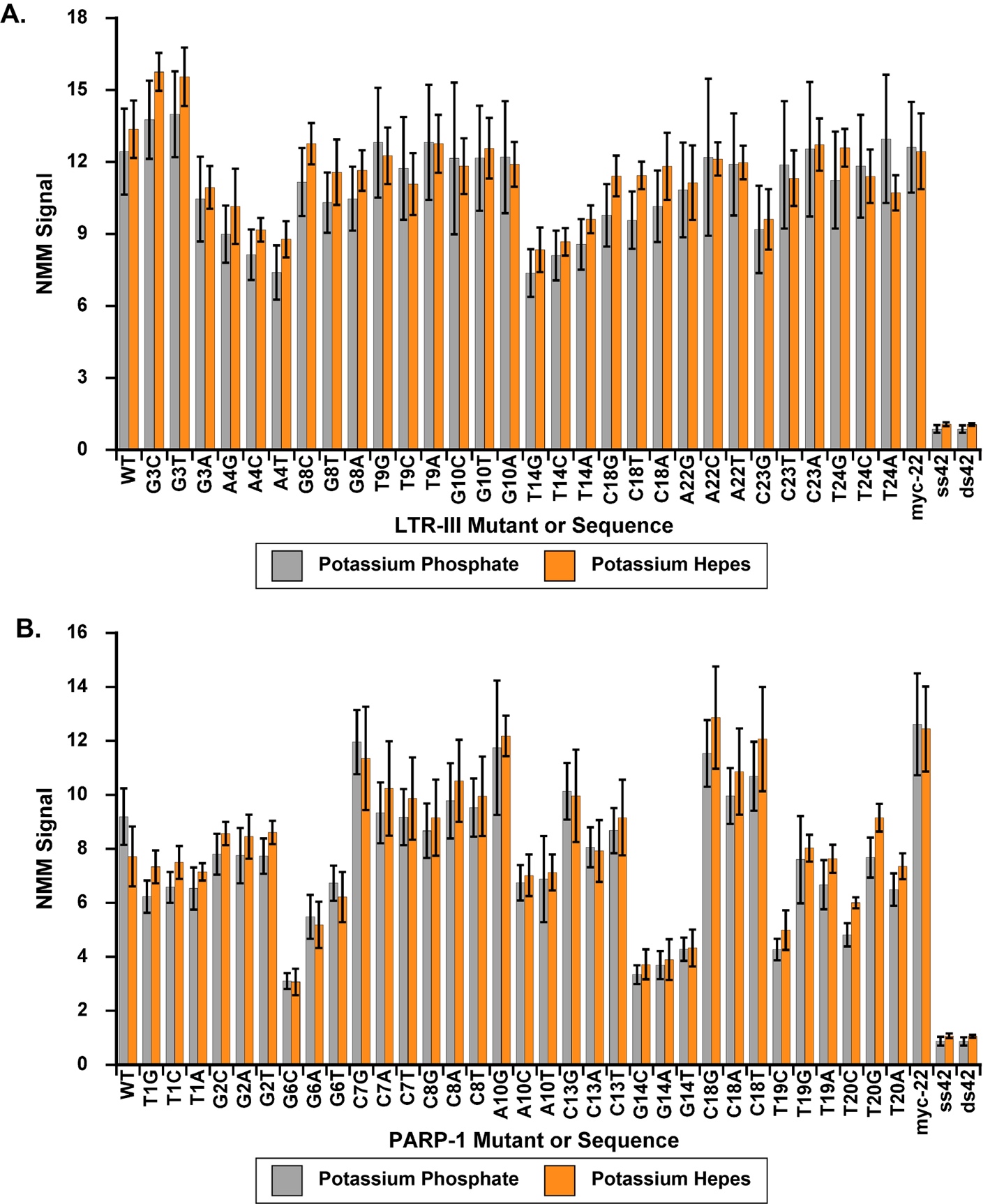

**SI Figure 2.** NMM turn-on ratio for **A.** LTR-III mutants and **B.** PARP-1 mutants in 10 mM potassium phosphate buffer pH 7.5 or 40 mM HEPES KOH buffer pH 7.5. The difference in buffers does not appear to affect G-quadruplex structure as NMM binding is consistent.

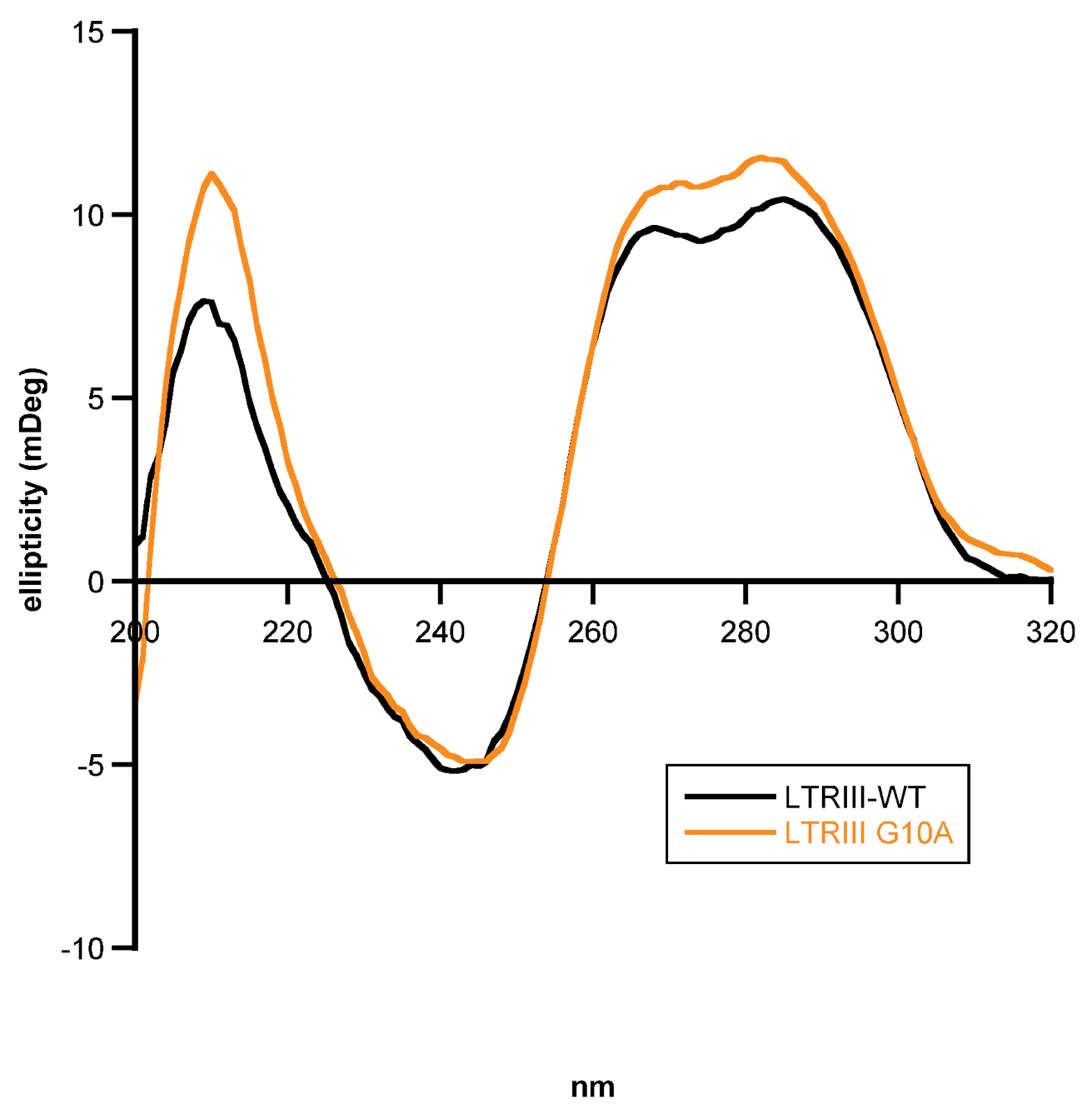

**SI Figure 3.** CD spectra comparing LTR-III WT and LTR-III G10A, overall the spectra are similar indicating the mutation didn’t change the topology despite large differences in activity.

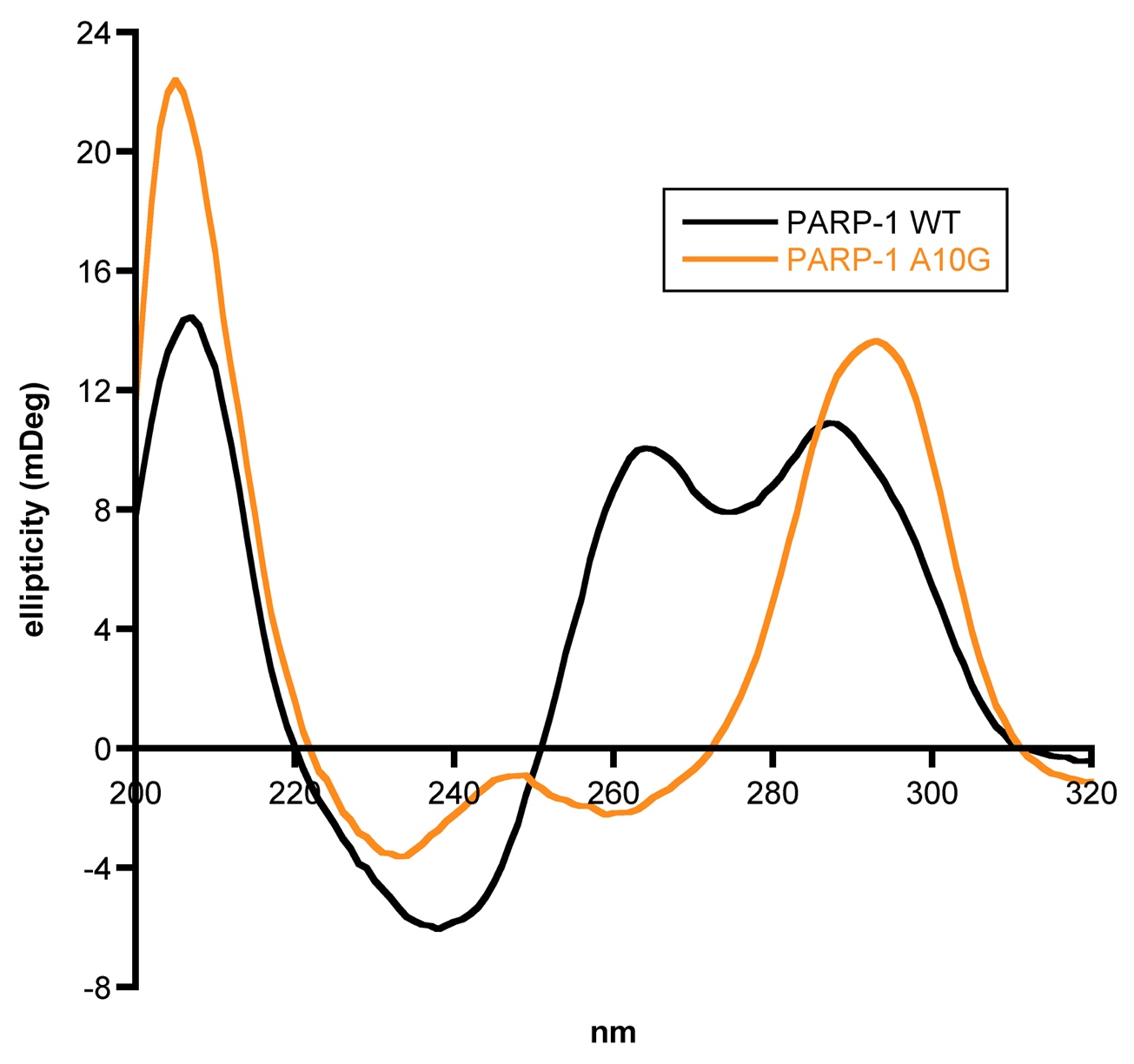

**SI Figure 4.** CD spectra comparing PARP-1 WT and PARP-1 G10A, the mutation results in the G-quadruplex adopting an anti-parallel topology under our conditions.

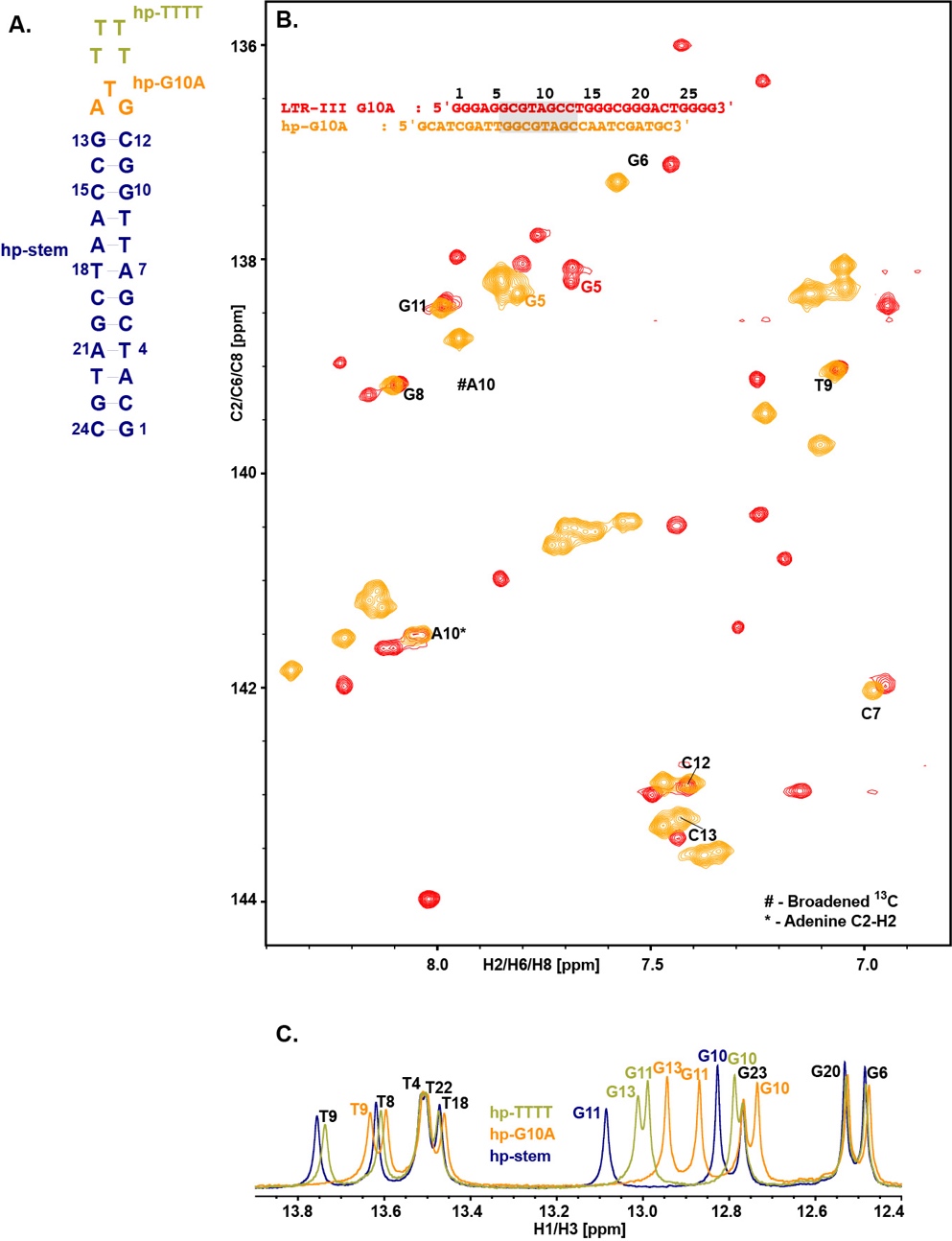

**SI Figure 5**. **A.** Sequence and predicted structure of the hairpin mutants hp-TTTT and hp-G10A. **B.** Overlay of ^13^C-^1^H region (aromatic) resonances of LTR-III G10A (red) and hp-G10A (orange), numbering indicates positioning according to LTR-III WT sequence. Closely matched resonances are marked in black and all other resonances in the sequence's respective colors. The # indicates broadened resonances in ^13^C dimension and * indicates ^13^C aliased resonances of adenine C2-H2 **C.** Overlay of ^1^H-1D imino resonances hp-TTTT (light green), hp-G10A (orange), and hp-stem (dark blue). The imino ^1^H peaks are indicative of hairpin and duplex DNA formation. Numbering indicates positioning according hp-stem sequence. Closely matched resonances are marked in black and all other resonances in the sequence’s respective colors.

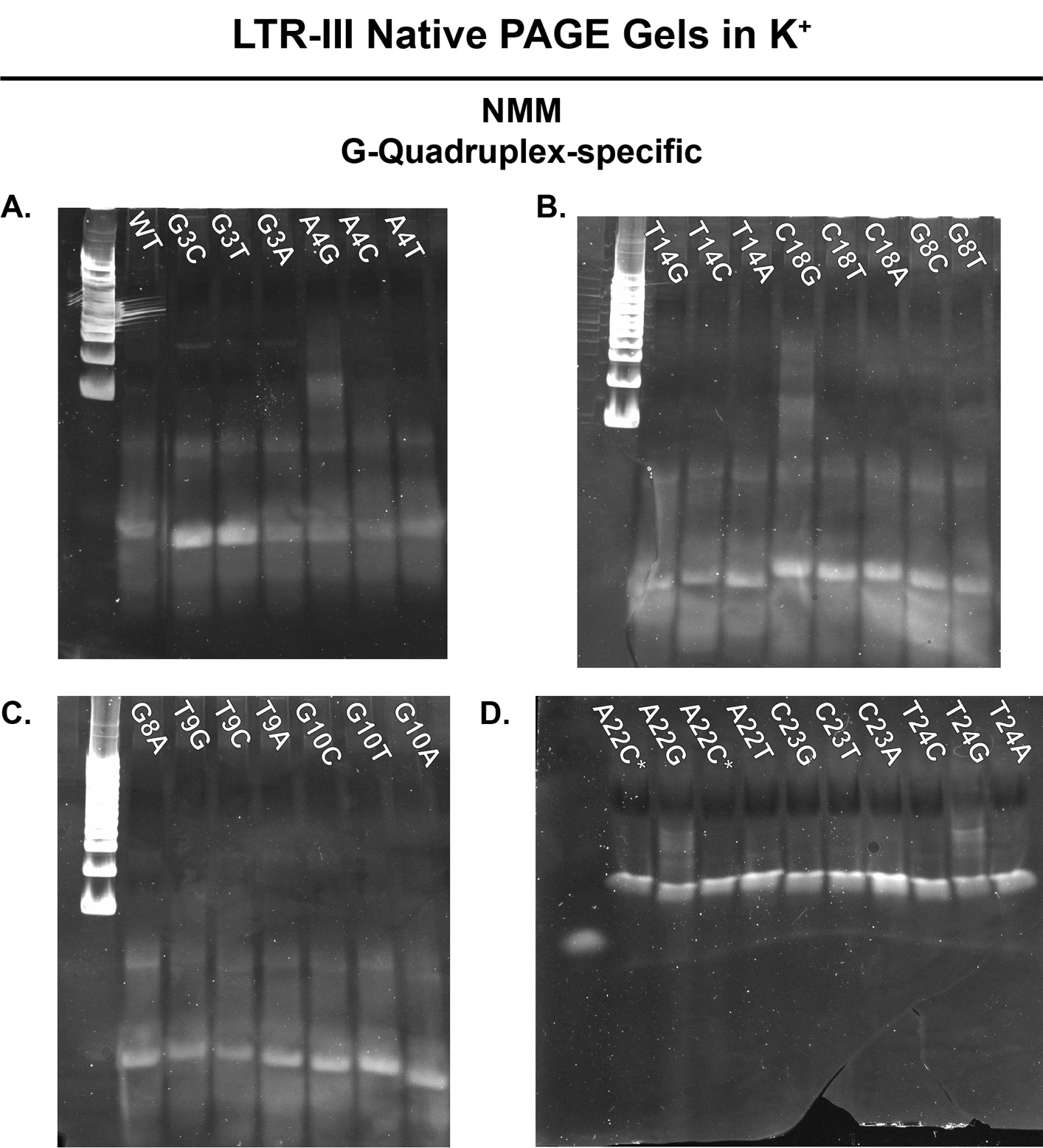

**SI Figure 6.** Non-denaturing native PAGE gels of LTR-III mutants in potassium containing TBE buffer, stained with NMM.

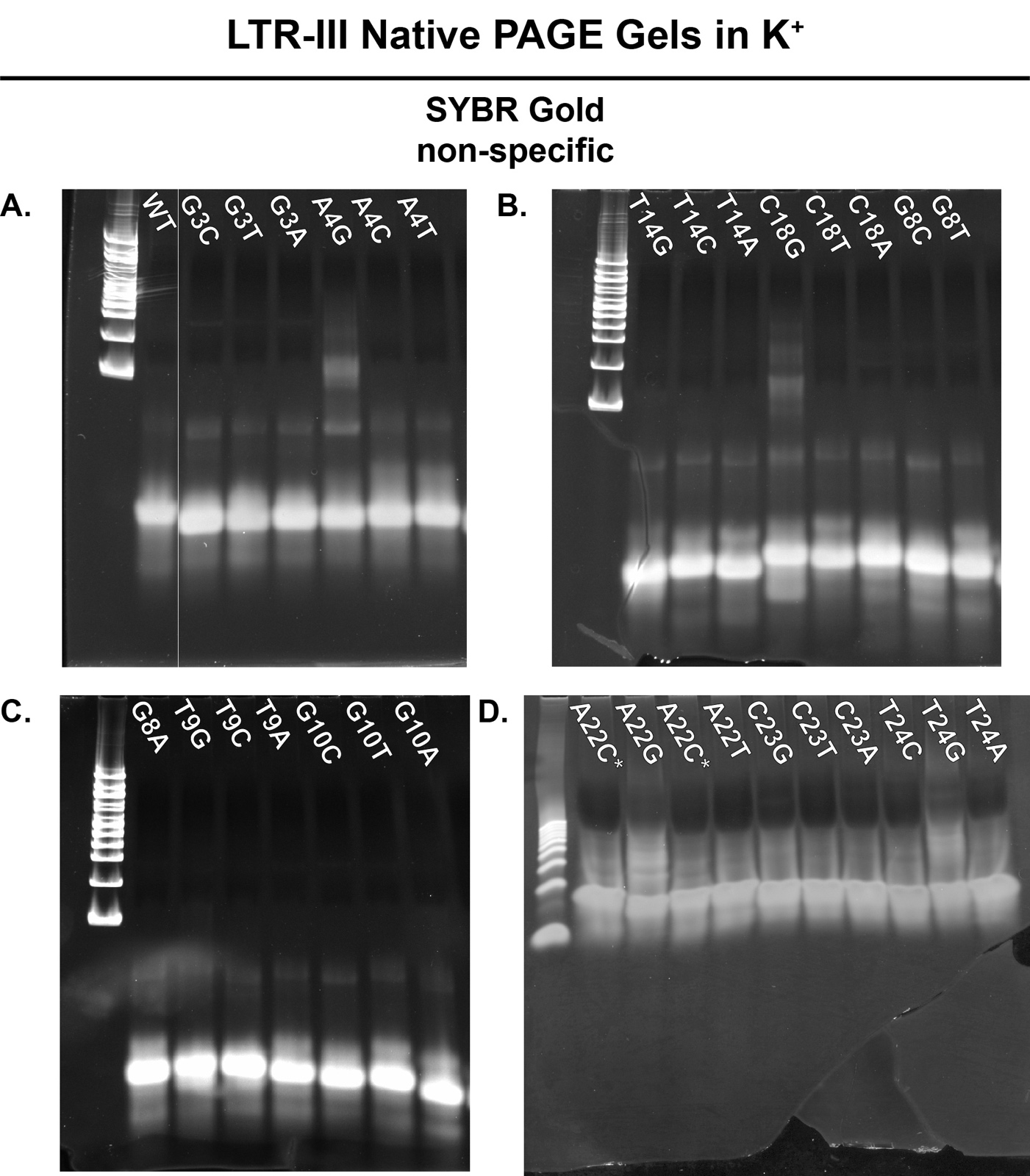

**SI Figure 7.** **A-D.** Non-denaturing native PAGE gels of LTR-III mutants in potassium containing TBE buffer, stained with SYBR gold. **A-C.** were for 1.5 hours, while D. was run for 1 hour.

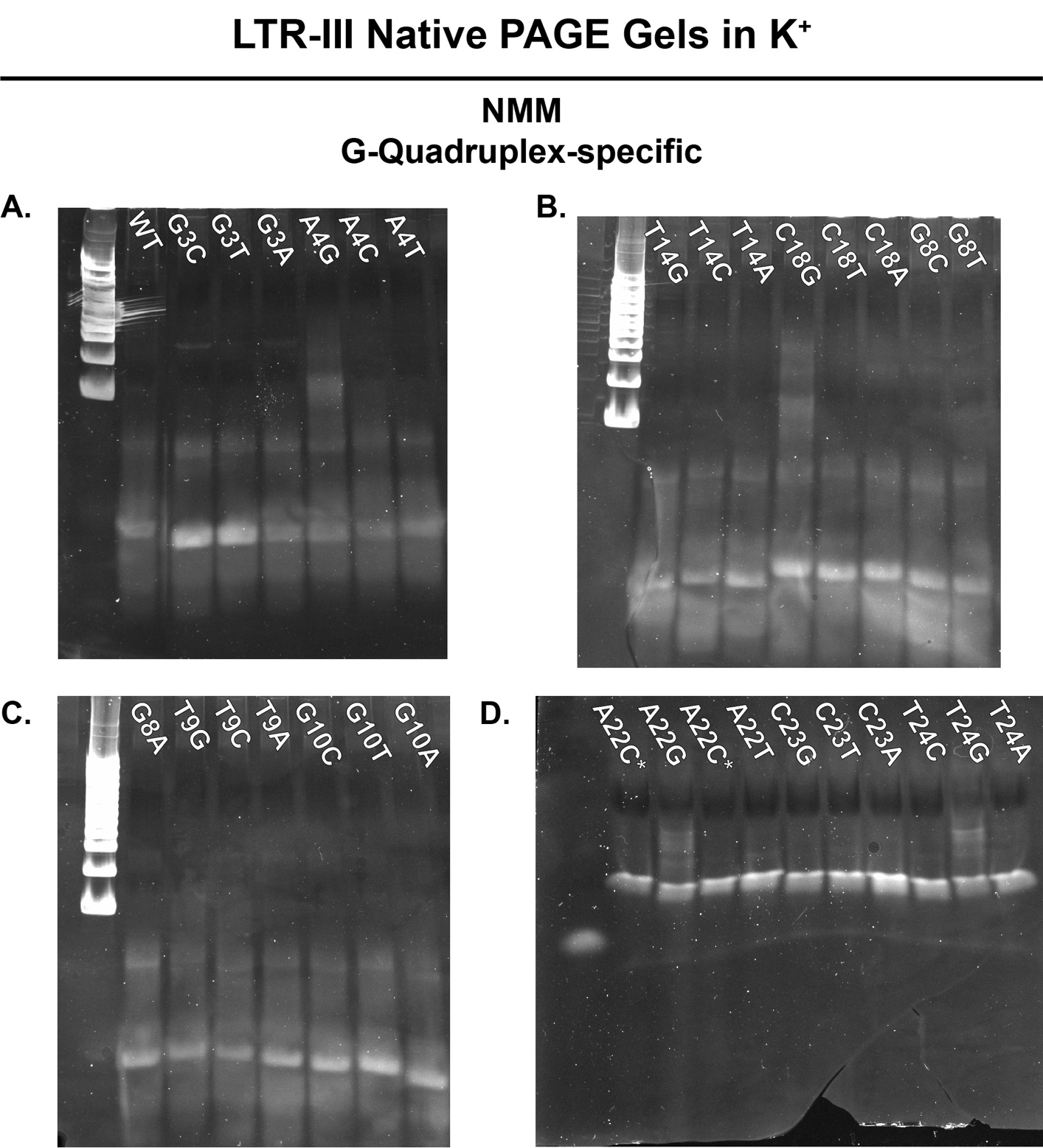

**SI Figure 8.** **A-D.** Non-denaturing native PAGE gels of LTR-III mutants in potassium containing TBE buffer, stained with NMM. **A-C.** were for 1.5 hours, while D. was run for 1 hour.

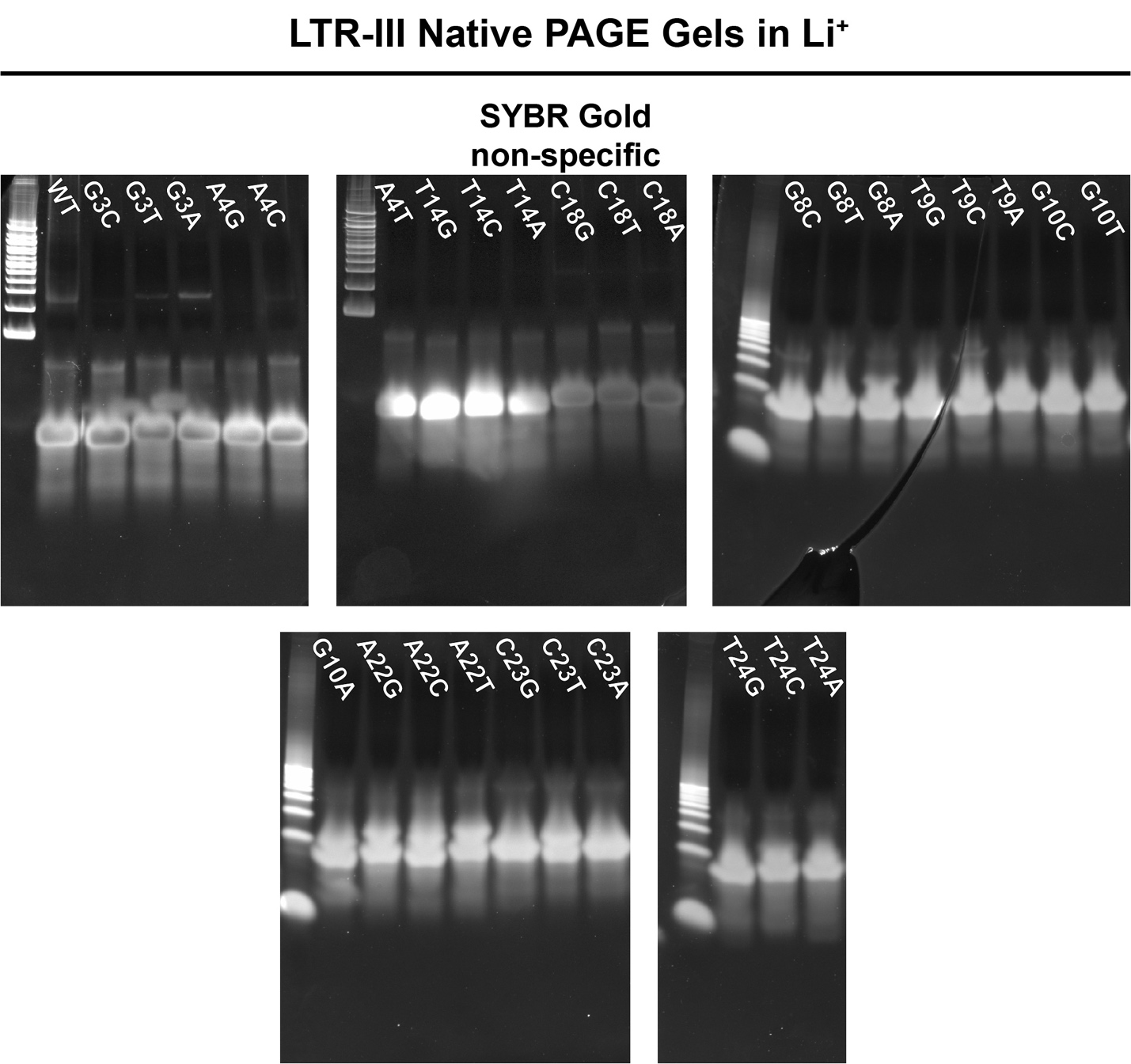

**SI Figure 9.** Non-denaturing native PAGE gels of LTR-III mutants in lithium containing TBE buffer, stained with SYBR gold.

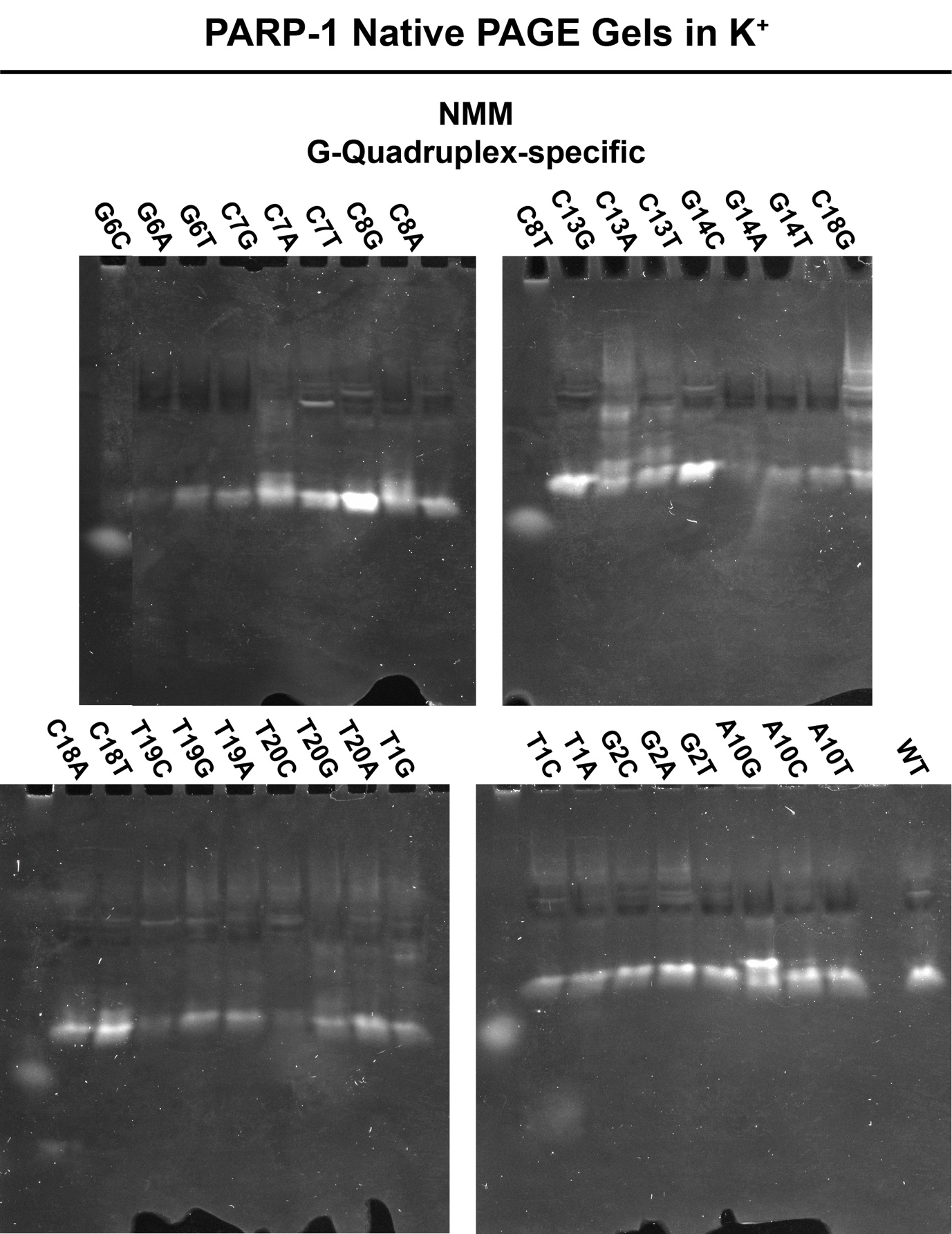

**SI Figure 10.** Non-denaturing native PAGE gels of PARP-1 mutants in potassium containing TBE buffer, stained with NMM.

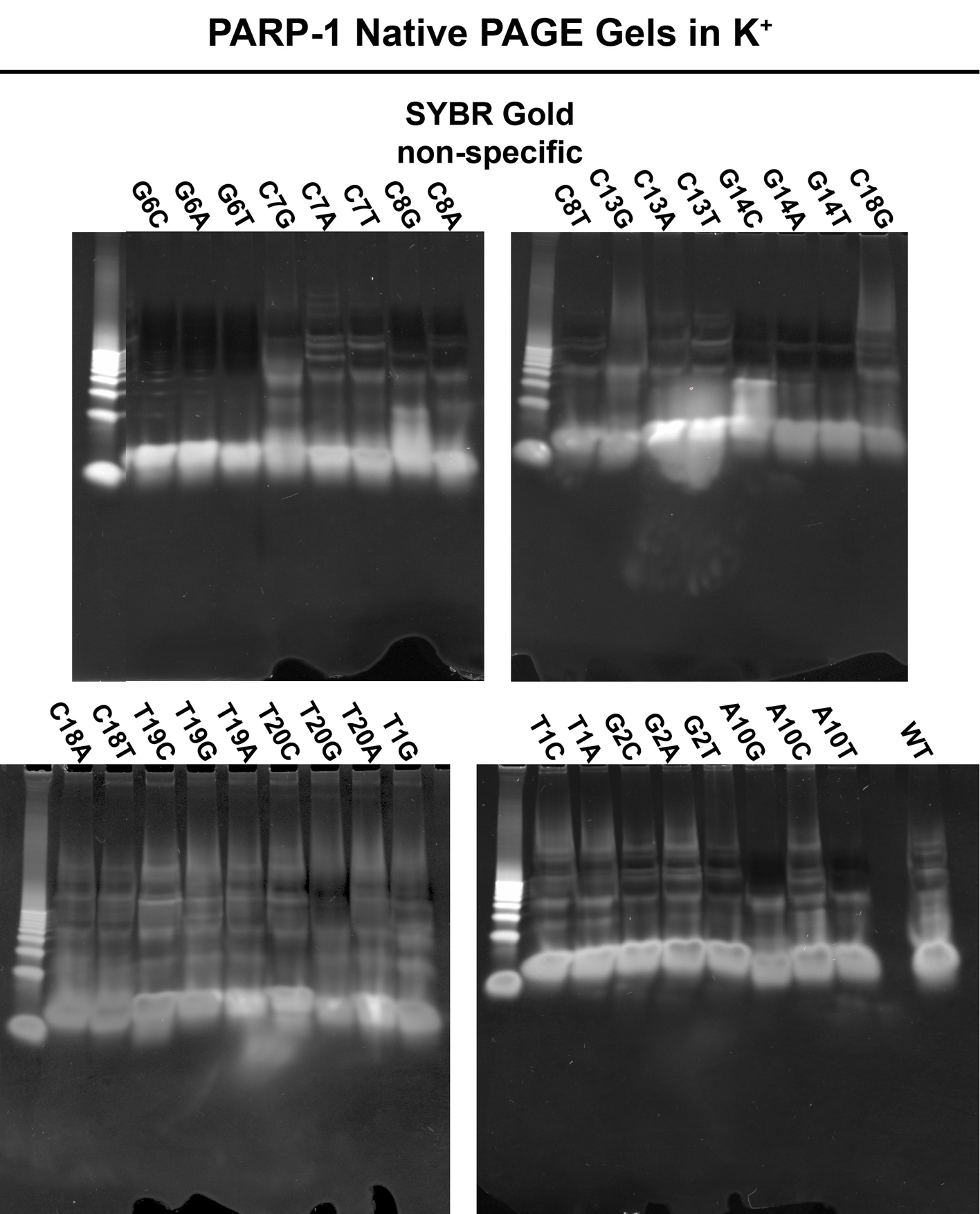

**SI Figure 11.** Non-denaturing native PAGE gels of PARP-1 mutants in potassium containing TBE buffer, stained with SYBR gold.

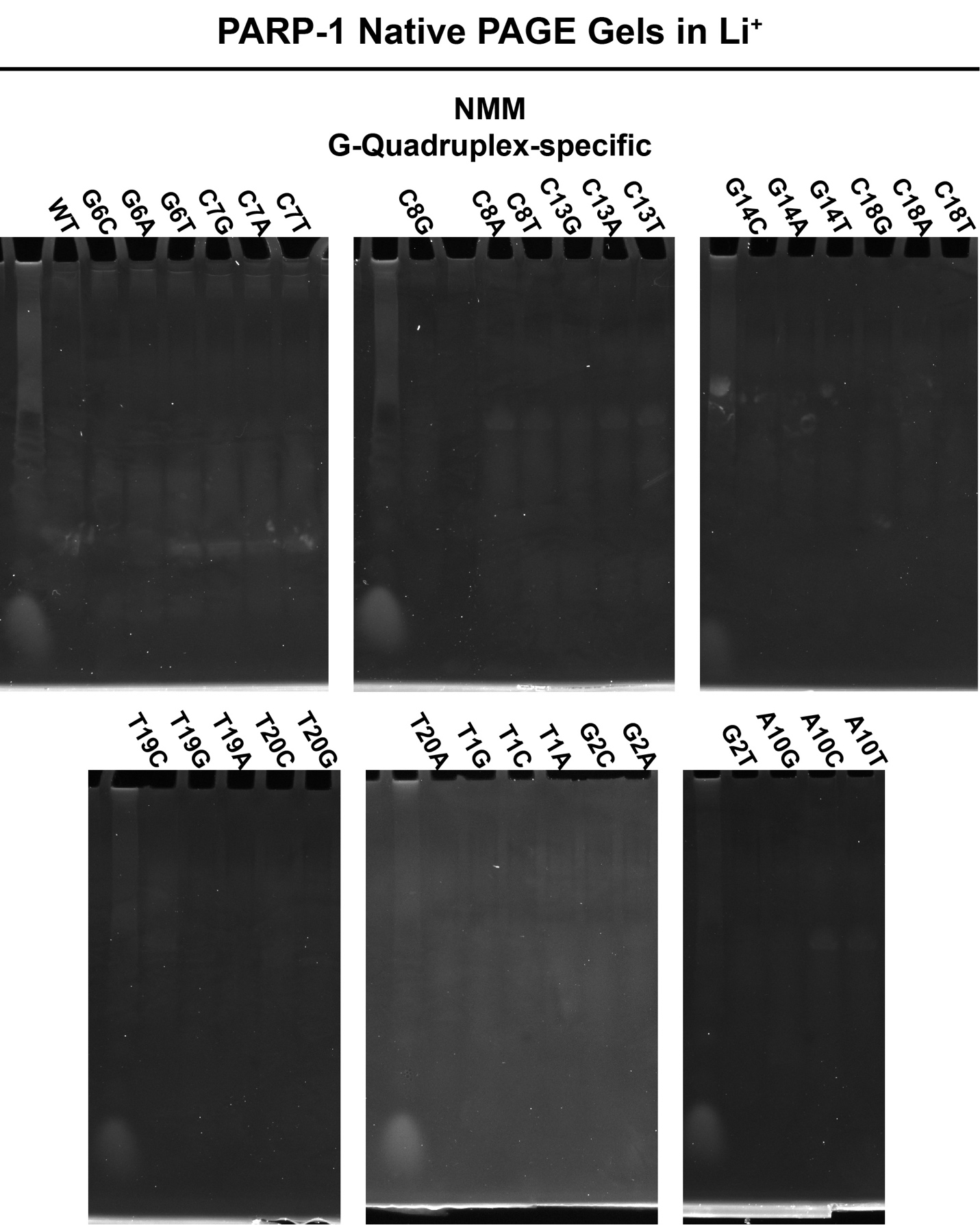

**SI Figure 12.** Non-denaturing native PAGE gels of PARP-1 mutants in lithium containing TBE buffer, stained with NMM.

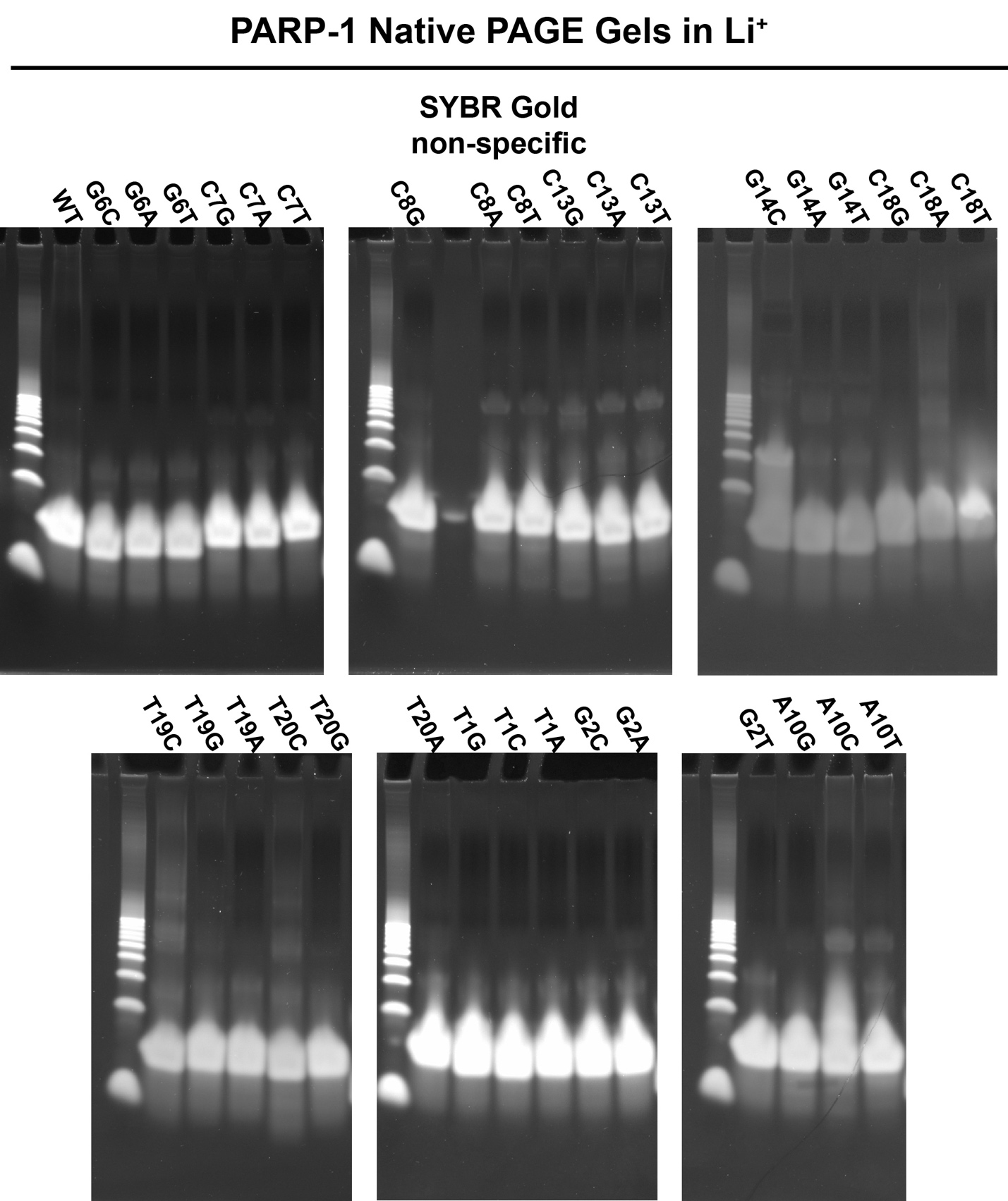

**SI Figure 13.** Non-denaturing native PAGE gels of PARP-1 mutants in lithium containing TBE buffer, stained with SYBR gold.
